# Random Walk with Restart on multilayer networks: from node prioritisation to supervised link prediction and beyond

**DOI:** 10.1101/2023.10.18.562848

**Authors:** Anthony Baptista, Galadriel Brière, Anaïs Baudot

## Abstract

**Background:** Biological networks have proven invaluable ability for representing biological knowledge. Multilayer networks, which gather different types of nodes and edges in multiplex, heterogeneous and bipartite networks, provide a natural way to integrate diverse and multi-scale data sources into a common framework. Recently, we developed MultiXrank, a Random Walk with Restart algorithm able to explore such multilayer networks. MultiXrank outputs scores reflecting the proximity between an initial set of seed node(s) and all the other nodes in the multilayer network. We illustrate here the versatility of bioinformatics tasks that can be performed using MultiXrank.

**Results:** We first show that MultiXrank can be used to prioritise genes and drugs of interest by exploring multilayer networks containing interactions between genes, drugs, and diseases. In a second study, we illustrate how MultiXrank scores can also be used in a supervised strategy to train a binary classifier to predict gene-disease associations. The classifier performance are validated using outdated and novel gene-disease association for training and evaluation, respectively. Finally, we show that MultiXrank scores can be used to compute diffusion profiles and use them as disease signatures. We computed the diffusion profiles of more than 100 immune diseases using a multilayer network that includes cell-type specific genomic information. The clustering of the immune disease diffusion profiles reveals shared shared phenotypic characteristics.

**Conclusion:** Overall, we illustrate here diverse applications of MultiXrank to showcase its versatility. We expect that this can lead to further and broader bioinformatics applications.

## Introduction

Random Walk is a powerful approach for analysing and exploring networks. By simulating the movement of a particle randomly traversing nodes and edges in a network, Random Walks are able to capture several topological and structural properties of networks [1], including connectivity [2], community structure [3], and node centrality [4]. Inspired from the PageRank algorithm [5], initially developed for ranking web pages in search results by simulating the behavior of an internet user following hyperlinks or restarting on arbitrary pages, Random Walk with Restart (RWR) was first introduced by Pan et al. [6]. In the RWR approach, the random particle, at each step, can navigate from one node to one of its neighbors or restart its walk from a node randomly sampled from a set of seed nodes. As PageRank, this strategy prevents the walker from getting trapped in dead ends and allows a more comprehensive exploration of the network’s topology [7]. RWR, by enabling restart from one or several seed nodes, simulates a diffusion process in which the objective is to determine the steady state of an initial probability distribution [8]. This steady state represents a measure of proximity between the seed(s) and all the network nodes, quantifying the extent to which the influence or information from the seed nodes has spread throughout the network. It overall identifies nodes that are closely connected to the seed(s) and provides valuable insights into the network’s organisation.

In computational biology, RWR has been particularly useful for the exploration of large-scale interaction networks and to derive guilt-by-association knowledge. For instance, RWR strategies significantly outperformed local distance measures for the prediction of gene-disease associations [9]. They have also been successfully applied to protein function prediction [10], identification of disease comorbidity [11], or drug-target interaction prediction [12]. More recently, RWR have been applied to drug prioritisation and repurposing for SARS-CoV-2 [13, 14].

Originally designed for investigating simple single-layer (i.e., monoplex) networks, RWR has been extended to navigate more complex networks, i.e. networks composed of multiple layers of interaction data. One such extension was proposed by Li and Patra [15] and introduced a RWR exploration of heterogeneous networks. They applied this approach to predict novel gene-phenotype relationships using a heterogeneous network composed of gene-gene interactions, phenotype-phenotype interactions, and known gene-phenotype associations. We introduced a RWR allowing the exploration of multiplex-heterogeneous networks, i.e., multiplex networks connected to each other by bipartite interactions [16]. More recently, we developed MultiXrank, a RWR algorithm able to explore generic multilayer networks [17]. We define a generic multilayer network as a multilayer network composed of any number and combination of multiplex and monoplex networks connected by bipartite interaction networks. In this multilayer framework, all the networks can also be weighted and/or directed. MultiXrank hence offers the opportunity to apply RWR on multilayer networks containing rich and complex interactions and fundamentally better suited for representing the multi-scale interactions observed in biological systems. In practice, MultiXrank outputs scores representing a measure of proximity between the seed(s) and all the nodes of the multilayer network. These output scores can then be used in a large number of downstream applications. We aim here to illustrate the versatility of the use of MultiXrank output scores. First, we show that MultiXrank can be used for node prioritisation. From a multilayer network containing gene, drug, and diseases interactions, we used MultiXrank scores to prioritise candidate drugs for leukemia. We also used the large network assembled in the Hetionet project [18], encompassing nine distinct types of nodes (including genes, drugs, diseases, biological processes, and pharmacological classes), to prioritise drugs for epilepsy. Second, we show that MultiXrank scores can be used to train a supervised classifier to predict gene-disease associations. Finally, we show how MultiXrank can be used to compute and compare diffusion profiles obtained for immune diseases on a multilayer network containing genomic information extracted from Promoter Capture Hi-C (PCHi-C) [19] experiments in different hematopoietic cells [20]. Overall, these diverse applications of MultiXrank demonstrate its versatility, both in the types of networks it can explore and the variety of downstream analyses that can be applied using its output scores.

## 1 Node prioritisation to study human genetic diseases

RWR approaches are frequently used to assess the proximity between seed node(s) and all the other nodes in a network. By leveraging the RWR output scores, nodes that are proximal to the seed node(s) can be prioritised. We will illustrate this prioritisation strategy by exploring the heterogeneous and rich information contained in biological multilayer networks using MultiXrank to prioritise genes and drugs in leukemia and epilepsy.

### 1.1 Prioritising genes and drugs of interest in Leukemia using MultiXrank on a gene and drug multilayer network

We first focused on leukemia, a disease for which we can confront our predictions with the knowledge accumulated in the literature. We prioritised genes and drugs of interest for leukemia based on MultiXrank output scores obtained from exploring a multilayer network composed of a gene multiplex network and a drug multiplex network, connected with a gene-drug bipartite network representing known drug-target associations (Materials and methods).

We selected two seeds associated with leukemia. More precisely, we selected HRAS as gene seed. HRAS is a gene of the RAS gene family associated with a wide variety of tumors, in particular in myeloid leukemia [21]. We also selected a drug seed, Tipifarnib (DB04960), a drug investigated for the treatment of acute myeloid leukemia and other types of cancer [22–24]. Using these two nodes jointly as seeds is particularly relevant as HRAS is a farnesylated protein and Tipifarnib is a farnesyltransferase inhibitor [25]. We applied MultiXrank (with the parameters specified in Supplementary Table S3) using these two seeds jointly and selected the top 10 highest-scoring gene and drug nodes (Supplementary Tables S4 and S5, respectively). We extracted the subnetwork connecting the seed nodes and the top 10 prioritised genes and drugs and their close neighborhood (Figure 1). We observed that prioritised nodes are close to both seeds, with a maximum shortest path distance between a prioritised node and a seed node equal to 4 (Supplementary Tables S4 and S5).

**Figure 1.**
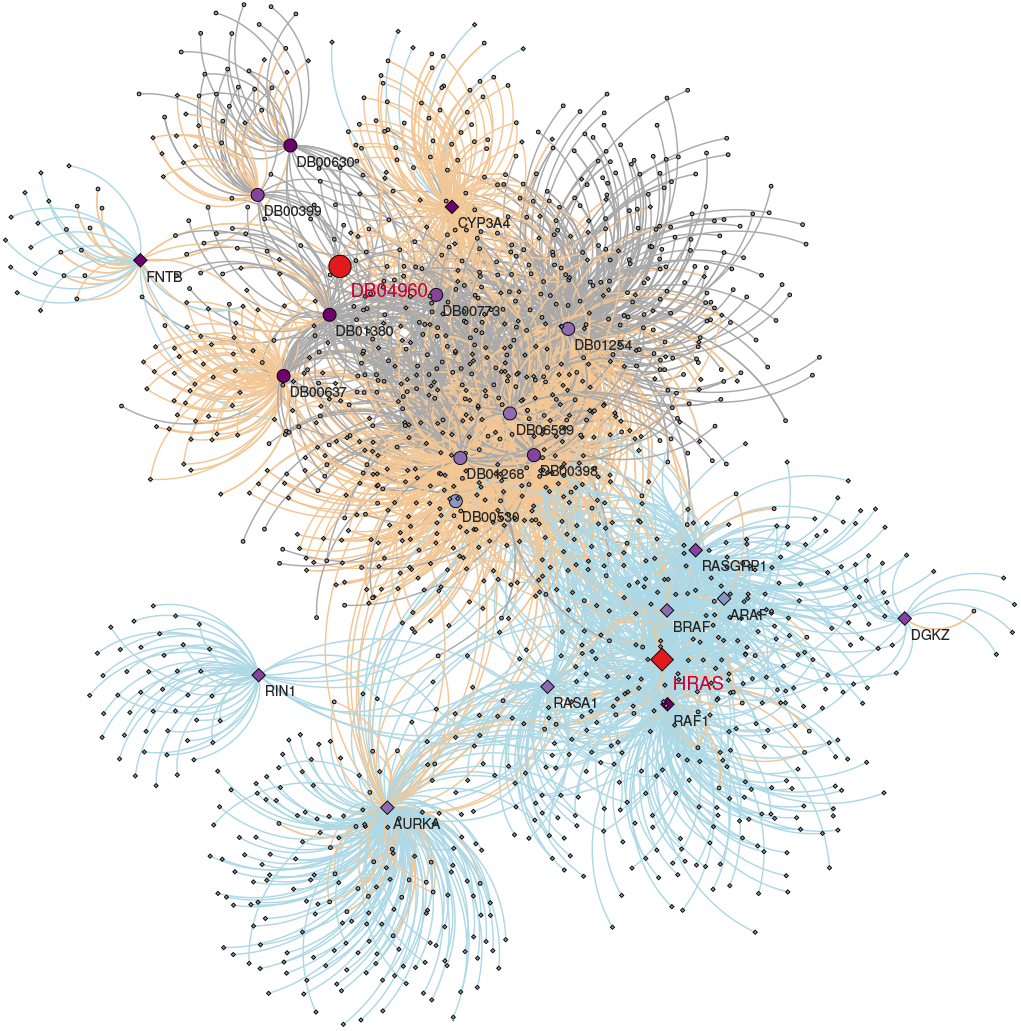
Subnetwork connecting the seed nodes (in red), the top 10 prioritised genes (diamonds) and drugs (dots) and their neighborhood. Vertex colors indicate the ranking of the nodes (except for the seed nodes, colored in red), with darker colors indicating better ranking. Edges are colored according to their provenance: gene multiplex (blue), drug multiplex (grey) and bipartite interactions (orange).

A literature survey of these top-10 prioritised drugs and genes establishes known or suspected connections with leukemia (Supplementary section 2.A). For instance, the top scoring gene, CYP3A4, is a drug-metabolising enzyme that has been shown to play a role in drug resistance in leukemia [26]. The second highest-scoring gene, FNTB, is coding the farnesyltransferase, and a target of Tipifarnib [27]. Different genes related to signal transduction and known to be relevant for cancer, such as RAF1, RASGRP1, RASA1, or ARAF, are also identified among the top-scoring genes. Moreover, the top prioritised drug, Astemizole (DB00637), is a good candidate for leukemia treatment as it’s anti-leukemic properties have been demonstrated in human leukemic cells [28]. Interestingly, Astemizole is metabolised by CYP3A4 [29], the top-scoring gene.

### 1.2 Prioritising genes and drugs of interest in Epilepsy using MultiXrank on a biomedical knowledge graph

We applied MultiXrank to prioritise candidate drugs for epilepsy, using as seed the epilepsy disease node (OID:1826) in the large and heterogeneous knowledge graph assembled in the Hetionet project [18]. This heterogeneous network is composed of eleven different types of nodes (Materials and Methods). We compared the drugs top-scored by MultiXrank, which is fully unsupervised, with the drugs prioritised by the Hetionet strategy, a supervised machine learning approach based on a regularised logistic regression model [18]. To evaluate the robustness of MultiXrank in relation to the choice of input parameters, we applied four distinct sets of parameters (Supplementary Table S6).

Most drugs are top-prioritised by both approaches. For instance, for one of the sets of parameters tested in MultiXrank (set of parameters number 4, Supplementary Table S6), 59% of the top-100 Hetionet prioritised drugs are also in the top-100 MultiXrank prioritised drugs, 79% are in the top-200 MultiXrank prioritised drugs, and 99% are in the top-500 MultiXrank prioritised drugs (Figure 2).

**Figure 2.**
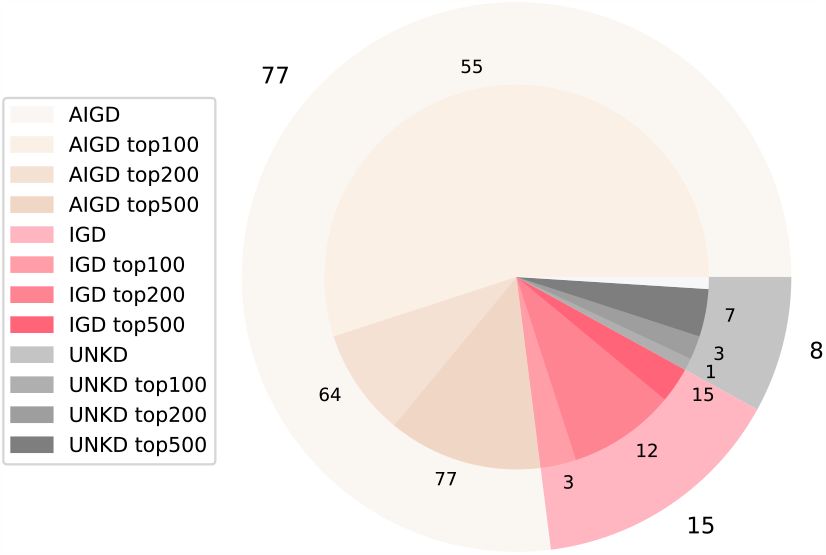
Ictogenic properties of the 100 drugs prioritised by Hetionet and overlap with the top 100, 200, and 500 drugs prioritised by MultiXrank (parameter set 4, see Supplementary Table S6). The data on ictogenic properties has been sourced from the Hetionet study [18]. AIGD are anti-ictogenic drugs that have a seizure suppressor effect (beige), IGD are ictogenic drugs (pink), and UKND are drugs with unknown effects (grey). The outer circle of the pie chart displays the number of AIGD, IGD, and UNKD drugs amongst the 100 drugs prioritised by Hetionet (e.g., 77 Hetionet prioritised drugs are AIGD). The center of the pie chart displays the number of drugs from each category that were also prioritised by MultiXrank in the top-100 (displayed in lighter shades), top-200 (displayed in middle shade), and top-500 drugs (displayed in darker shades). For instance, 55 top-100 MultiXrank prioritised drugs are AIGD and 64 top-200 MultiXrank prioritised drugs are AIGD. The drugs prioritised in the top-500 include the drugs prioritised in the top-200, and the top-200 includes the drugs prioritised in the top-100. The white part of the pie chart corresponds to the drugs prioritised by Hetionet that are not in our MultiXrank top-500 prioritised drugs.

We further checked the 41 drugs from the top-100 drugs identified by MultiXrank that are not prioritised by Hetionet (Supplementary Table S7). Interestingly, 3 of them (namely, Propofol, Vigabatrin and Diclofenac, respectively ranked 8, 23 and 49 by MultiXrank) have been tested in clinical trials for epilepsy, according to the DrugBank [30]. After extracting the DrugBank Categories associated to those 41 drugs (Supplementary Table S7), we observed that 24 of them are classified as *Cytochrome P-450 Substrates*.A recent study has shown that spontaneous recurrent seizures in mice modify Cytochrome P-450 expression in the liver and hippocampus. The authors hypothesise that nuclear receptors or inflammatory pathways can be considered as candidates for Cytochrome P-450 regulation during seizures [31]. Another study showed that Cytochrome P-450 enzymes can have a significant impact on the response to anti-epileptic drugs [32]. The second most represented DrugBank Category in the list of the 41 drugs prioritised by MultiXrank but not by Hetionet was the category *Agents that produce hypertension*, which map to 18 drugs. A review of the existing literature regarding hypertension and epilepsy show that those two conditions often co-occur [33, 34]. Furthermore, the relationship between the two conditions could be bidirectional, meaning that they can influence and exacerbate each other [35].

These results indicate that MultiXrank can provide predictions complementary to the Hetionet supervised machine learning approach. In addition, MultiXrank predictions can be easily interpreted as the subnetworks underlying the top-scoring nodes can be easily extracted.

## 2 Supervised prediction of gene-disease associations

In a second study, we present a supervised approach to predict gene-disease associations. Predicting gene-disease associations is crucial for the diagnosis, understanding, and treatment of genetic diseases. Among available approaches to predict gene-disease associations, network-based methods have been particularly exploited and have demonstrated good performances [36]. These network approaches were initially based mainly on unsupervised strategies, but an increasing number of methods are implementing supervised strategies [36]. Here, we use the output scores of MultiXrank to train supervised XGBoost and Random Forest binary classifiers to predict gene-disease associations (Supplementary Figure S5).

We used a multilayer network composed of a gene multiplex network and a disease monoplex network (Materials and Methods). These multiplex and monoplex networks are connected by a gene-disease bipartite network constructed with an outdated version of DisGeNET (v2.0, 2014, [37]). The edges of the bipartite network are weighted according to the support score provided by DisGeNET v2.0 (2014). We applied MultiXrank on the multilayer network described above, using the gene and disease nodes from each gene-disease association as seeds. The parameters used for running MultiXrank are detailed in Supplementary Table S8. We used both positive associations (i.e. true gene-disease associations) and negative associations (i.e., random gene-disease pairs that are not associated according to DisGeNET). For each set of positive seeds (true gene-disease association), the gene-disease bipartite edge connecting the two seeds was removed from the bipartite network before training.We collected MultiXrank output scores obtained for all positive and negative gene-disease pairs of seeds and trained binary XGBoost and Random Forest classifiers with different parameters (Supplementary Table S9). We then tested the performance of the classifiers in predicting unseen gene-disease associations from the outdated version of DisGeNET that were kept out for testing as well as the gene-disease associations that have been added in the updated version of DisGeNET (v7.0, 2020, [38]). The full machine learning procedure is detailed in the Materials and Methods section. We also report the performances of our models in predicting DisGeNET v2.0 (2014) and DisGeNET v7.0 (2020) associations in Supplementary Tables S9 and S10, respectively. For the prediction of unseen test DisGeNET v2.0 (2014) associations, the best classification performance was achieved with an XG-Boost model taking class imbalance into account. This model reached a balanced accuracy of 0,85 and an F1-score of 0,79, showing the predictive potential of MultiXrank output scores. However, the prediction performance dropped considerably for predicting DisGeNET v7.0 (2020) associations (balanced accuracy 0,64 and F1-score 0,53). It should be noted that the MultiXrank scores used for the classification of DisGeNET v2.0 (2014) and DisGeNET v7.0 (2020) associations were calculated on the same network, constructed solely from the information contained in DisGeNET v2.0 (2014). Importantly, DisGeNET v2.0 (2014) reported only 381 654 gene-disease associations, whereas DisGeNET v7.0 (2020) reported 1 135 037 associations, which represents a threefold increase. Moreover, over the 21 666 genes reported in DisGeNET v7.0 (2020), only 14 255 appeared in DisGeNET v2.0 (2014). Similarly, only 38% of the 30 170 diseases reported in DisGeNET v7.0 (2020) were also reported in DisGeNET v2.0 (2014). The substantial increase in the amount of information contained in DisGeNET between 2014 and 2020 is potentially the cause of the significant decrease in classification performance.

## 3 Diffusion profiles comparison to unveil immune diseases similarities

The scores resulting from a random walk using a given seed can be regarded as a diffusion profile and represent a network-based molecular signature. Diffusion profiles obtained starting from different seeds can then be compared to reveal signature proximities. Here, we propose to compute and compare the diffusion profiles obtained using 131 immune diseases (Supplementary Table S13) as seed in MultiXrank applied to several multilayer networks. We created eight hematopoietic cell-specific multilayer networks, each composed of four different monoplex/multiplex networks (Figure 3, Materials and Methods). The first two monoplex/multiplex networks incorporate gene and disease interactions sourced from public databases. These two layers are the sames across all the eight multilayer networks. The remaining two layers encode Promoter Capture Hi-C (PCHi-C) fragment interactions and Topologically Associating Domain (TAD) interactions observed in various hematopoietic cell lines, extracted from a dataset generated in [20]. It’s important to note that the PCHI-C fragment layer and the TAD layer are unique to each hematopoietic cell line, and hence vary across the eight multilayer networks (Supplementary Section 1.C).

**Figure 3.**
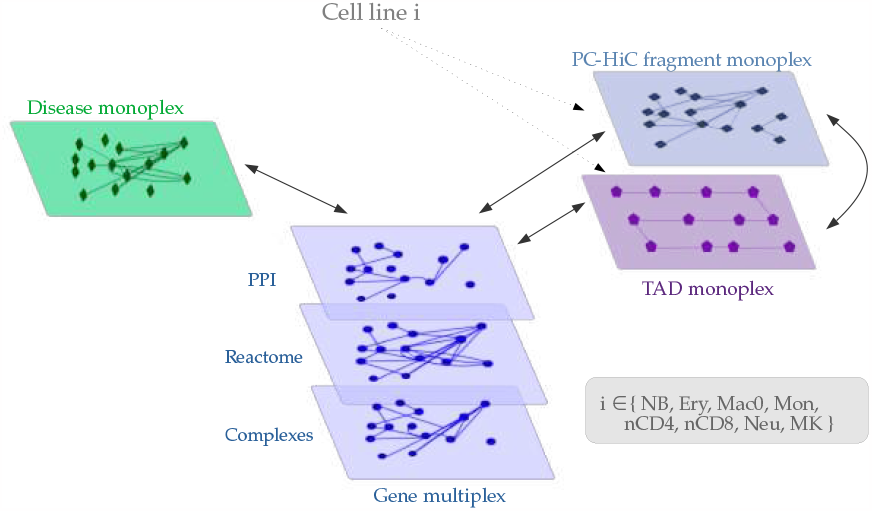
Hematopoietic multilayer networks composed of two genomic layers built from PCHi-C and TAD data, of a gene multiplex network and of a disease monoplex network. The disease monoplex and the gene multiplex network are the same in all the hematopoietic multilayer networks. However, the PCHi-C and TAD layers are specific to each hematopoietic cell line. The black arrows represent the bipartite networks that connect two different types of nodes.

The strength of this multilayer network constructions lies in its capacity to combine non cell-specific generic gene and disease interactions with data regarding genomic interactions unique to hematopoietic cell lineages. In addition, the genomic interaction layers allow us to consider data representing the 3D conformation of DNA and non-coding regions of the genomes. This 3D conformation of DNA is a key to understanding, for instance, genomic structural variations that are key players in the study of diseases [39]. We demonstrated that these genomic data maintain the signal of the hematopoietic cell type. Indeed, the PCHi-C fragments and TAD datasets capture the tree lineage of hematopoietic cells (Supplementary Section 4.A, Supplementary Figure S6). We also demonstrate that this lineage signal is captured in the RWR scores obtained from applying MultiXrank to the eight multilayer networks (Supplementary Section 4.B, Supplementary Figures S7 and S8).

Here, we aim to apply MultiXrank on the eight hematopoietic multilayer networks using as seeds 131 different immune diseases to obtain the disease diffusion profiles. We consider the diffusion profiles as disease signatures. We will next cluster the immune diseases based on the similarity of their diffusion profiles. We hypothesise that such clustering can reveal potentially similar immune diseases.

To reveal similarities between the 131 immune diseases based on the diffusion profiles obtained on the eight multilayer networks, we first compute disease-disease distances for each cell type (i.e. hematopoietic multilayer network) and node type (i.e. disease nodes, protein nodes, PCHi-C fragment nodes and TAD nodes) (equation 1, Materials and Methods).

Then, the disease-disease distance matrices obtained for the eight hematopoietic multilayer networks are fused (equation 2, Materials and Methods). This procedure produces 4 disease-disease integrated similarity matrices, one for each node type in the multilayer networks. These matrices are then used to cluster the immune diseases using a multiview clustering algorithm (Materials and Methods). The obtained clusters are detailed in Supplementary Table S12. Additionally, the 4 matrices are concatenated into a single matrix and projected into a 2D t-SNE (t-distributed stochastic neighbor embedding) [40] space. In this projection, we label each immune disease with their corresponding cluster (Figure 4).

**Figure 4.**
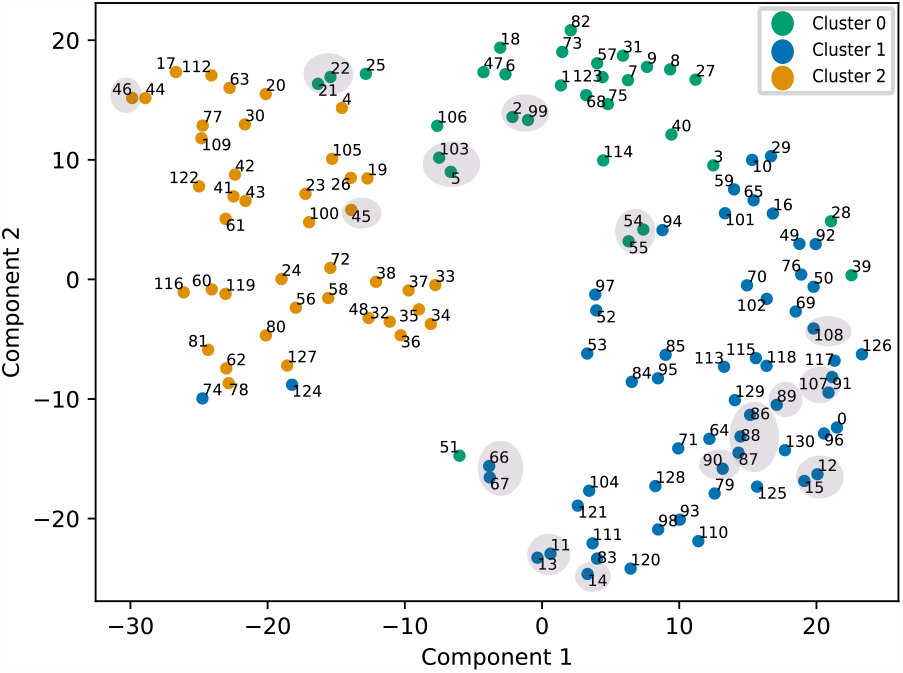
t-SNE projection of the integrated distances between the 131 different immune diseases. Colors indicate the clusters in which the diseases were grouped according to the multiview spectral clustering algorithm. Points highlighted with grey background correspond to diseases cited in the main text.

Interestingly, we can assess the relevance of the approach by examining diseases that represent distinct subtypes of the same condition. These disease subtypes are caused by different mutated genes and hence present different connection patterns in the gene and disease monoplex/multiplex networks. Nevertheless, being subtypes of the same condition, we expect these diseases to have similar network-based diffusion profiles and cluster together. Our examination of the results substantiates this, as demonstrated for all diseases subtypes included in our list of 131 immune diseases, listed below:

- Autosomal recessive early-onset inflammatory bowel disease 28 (UMLS:C2751053, seed 54) and autosomal recessive early-onset inflammatory bowel disease 25 (UMLS:C2675508, seed 55), both grouped in cluster 0 and close in the t-SNE space (Figure 4, cluster 0) ;
- Hypogammaglobulinemia AGM2 (UMLS:C3150750, seed 11), hypogammaglobulinemia AGM3 (UMLS:UMLS:C3150751, seed 12), hypogammaglobulinemia AGM4 (UMLS:C3150752, seed 13), hypogammaglobulinemia AGM5 (UMLS:C3150753, seed 14) and hypogammaglobulinemia AGM6 (UMLS:C3150207, seed 15), all grouped in cluster 1. In the t-SNE space, hypogammaglobulinemia AGM2, AGM4 and AGM5 are close, as well as hypogammaglobulinemia AGM3 and AGM6 (Figure 4, cluster 1) ;
- Immunodeficiency with hyper IgM type 1 to type 5 (UMLS:C0398689 (seed 86), UMLS:C1720956 (seed 87), UMLS:C1720957 (seed 88), UMLS:C1842413 (seed 89), UMLS:C1720958 (seed 90)), all grouped in cluster 1 and close in the t-SNE space (Figure 4, cluster 1) ;
- complement component 8 deficiency type 1 (UMLS:C3151081, seed 66) and Complement component 8 deficiency type 2 (UMLS:C3151080, seed 67), both grouped in cluster 1 and close in the t-SNE space (Figure 4, cluster 1) ;
- Activated PI3K-Delta Syndrome 1 (UMLS:C3714976, seed 108) and activated PI3K-Delta Syndrome 2 (UMLS:C4014934, seed 107) are grouped in cluster 1 and close in the t-SNE projection (see Figure 4, cluster 1).
- Aicardi-Goutières syndrome 1 (UMLS:C0796126, seed 45) and AicardiGoutières syndrome 2 (UMLS:C3489724, seed 46) are grouped in cluster 2 (Figure 4, cluster 2).

We conducted additional analysis to explore the composition of disease clusters and extract their essential characteristics:

- **Cluster 0** regroups 31 immune diseases. It is mainly composed of Inflammatory (e.g. Takayasu’s arteritis, giant cell arteritis (temporal arteritis), autosomal recessive early-onset inflammatory bowel disease), Autoinflammatory (e.g. tumor necrosis factor receptor-associated periodic syndrome (TRAPS), hyper-IgD syndrome, familial cold autoinflammatory syndrome), Autoimmune diseases (e.g: rheumatoid arthritis, type 1 diabetes, cicatricial pemphigoid, Hashimoto’s thyroiditis). This grouping underscores the intricate relationship between inflammation, auto-inflammation and autoimmunity, as supported by the existing literature on inflammatory disorders. Indeed, numerous studies have postulated an immunological continuum linking monogenic autoinflammatory disorders with autoimmunity [41, 42].
- **Cluster 1** encompasses 59 immune diseases. It groups conditions marked by immunodeficiencies, primary immunodeficiencies (including several types of Complement component deficiencies) and increased susceptibility to infections. Within this cluster, numerous diseases are linked to immunoglobulin-related abnormalities, including various forms of hypogammaglobulinemia, immunodeficiency with hyper IgM, as well as immunoglobulin A deficiency and agammaglobulinemia.
- **Cluster 2** accounts for 41 immune diseases. These diseases appear to be rather diverse, encompassing diseases that impact a wide range of bodily systems. Many types of leukemias (e.g. acute myeloid leukemia, chronic lymphocytic leukemia, chronic myelogenous leukemia), lymphomas (e.g. Hodgkin’s lymphoma, non-Hodgkin lymphoma) and other blood-related diseases (e.g. pernicious anemia, myelodysplastic syndromes) are included in this cluster. Other systems affected by diseases from cluster 2 include the cardiovascular (e.g: congenital heart block), hepatic (e.g. glycogen storage disease type 1B), skeletal (e.g. cherubism), dermatological (e.g. lichen sclerosus, pruritic urticarial papules plaques of pregnancy), muscular (e.g. inclusion body myositis), neurological (e.g. Aicardi-Goutieres syndrome) and neuromuscular (e.g. stiff person syndrome) systems. However, many of these diseases can impact multiple systems (e.g. DiGeorge syndrome, Aicardi-Goutières syndrome, CHARGE syndrome, glycogen storage disease type 1B, Pearson syndrome).

Facing the apparent diversity of diseases included in cluster 2, comparatively to the well defined clusters 0 and 1, we further investigated the composition of cluster 2. A literature review on the diseases included in this cluster shows that most of them are associated with blood diseases on one hand and cardiovascular system diseases on the other hand. Indeed, many of the diseases that are not directly associated with leukemia or lymphoma appear to be comorbid to those cancers. For instance, elevated risks of lymphoma and leukemia have been reported for patients with pernicious anemia [43] and myelodysplastic syndromes can evolve to acute myeloid leukemia [44]. Other diseases from cluster 2 that are considered associated with lymphoma and leukemia in the literature include ataxia telangiectasia [45], Bloom syndrome [46], cartilage-hair hypoplasia [47], and Chediak-Higashi syndrome [48], among others. Moreover, many diseases from cluster 2 could be associated with cardiovascular diseases, according to the literature: congenital heart defects are observed in 50–85% of cases of CHARGE syndromes [49] ; the TARP syndrome is associated with congenital heart defects [50] ; congenital heart disease is a common feature in the 22q11.2 deletion syndrome (DiGeorge syndrome) [51] ; Parry Romberg syndrome is associated with hypertrophic cardiomyopathy and rheumatologic heart disease [52] ; lichen sclerosus is associated with increased risk of cardiovascular comorbidities in female [53] ; Pearson syndrome is often associated with cardiac conduction defects [54] ; the association of inclusion body myositis and cardiac disease is debated [55] ; a case of Melkersson–Rosenthal syndrome affecting cardiac connective tissues was reported in [56].

Finally, we examined some diseases close to each other in the t-SNE space. We detail here 3 examples:

- Chronic recurrent multifocal osteomyelitis (CRMO) (UMLS:C0410422, seed 2) and Majeed syndrome (UMLS:C1864997, seed 99) (Figure 4, cluster 0): CRMO is known as one of the major features of Majeed syndrome [57].
- Netherton syndrome (UMLS:C0265962, seed 103) and Eosinophilic esophagitis (UMLS:C0341106, seed 5): (see Figure 4, cluster 0): Eosinophilic esophagitis is observed in 44% of the people with Netherton syndrome [58].
- Myasthenia gravis (UMLS:C0026896, seed 21) and Myositis (UMLS:C0027121, seed 22) (see Figure 4, cluster 0): several cases of co-existence of Myasthenia Gravis and Myositis are reported in [59].

All of these observations suggest that the integrated MultiXrank scores effectively capture similarities in disease diffusion profiles, indicating shared phenotypic manifestations and potential comorbidity patterns among the diseases.

## Conclusion

Multilayer networks provide a valuable framework for integrating a wide range of biological interactions involving diverse types of entities. The MultiXrank Random Walk with Restart algorithm can effectively explore such multilayer networks, offering opportunities for various analyses. In this study, we demonstrate the versatility of the MultiXrank algorithm through three distinct biological applications: prioritising drugs and genes in a disease context, predicting gene-disease associations, and comparing and clustering diseases.

## 4 Materials and methods

### 4.1 Construction of the biological networks

The different studies presented in this manuscript explore different biological networks summarised in Supplementary Table S1. Detailed information on the biological networks used in each study are provided in Supplementary Section 1.

In study 1, for node prioritisation in leukemia, we used a multilayer network composed of a gene and a drug multiplex networks, connected by bipartite gene-drug interactions (Supplementary Section 1.A, Supplementary Table S1). In the gene multiplex network, we encode 3 types of gene interactions: protein-protein interactions, molecular complexes and pathways gathered from public databases. In the drug multiplex network, we encode 4 types of drug interactions: adverse interactions, experimental drug combinations, computationally predicted drug interactions and pharmacological interactions. The two multiplex networksare connected with gene-drug associations extracted from the repoDB database [60].

In study 1, for node prioritisation in epilepsy, we used Hetionet [18], an network containing 11 types of nodes (Supplementary Section 1.B, Supplementary Table S1). The first multiplex network encodes 3 types of gene interactions: coexpression, physical interaction and regulation. The second and third monoplex networks encode for disease similarities and drug similarities, respectively. The three monoplex/multiplex networks are connected with bipartite gene-drug interactions, gene-disease interactions and disease-drug interactions. The multilayer network also contains other node types (including pathways, biological process and pharmacologic classes) that are connected to the gene, drug and disease networks via bipartite interactions.

In study 2, Gene-Disease associations were predicted from MultiXrank output scores obtained by exploring a multilayer network composed of a gene multiplex network and a disease monoplex network, connected by gene-disease bipartite interactions (Supplementary Section 1.A, Supplementary Table S1). The multiplex network encodes gene interactions, identical to the one used in node prioritisation for leukemia. The monoplex network encode disease interactions. The gene-disease bipartite associations were extracted from an outdated version of the DisGeNET database (v2.0, 2014).

Finally, in study 3, we obtained MultiXrank scores from the exploration of eight hematopoietic multilayer networks (Supplementary Section 1.C, Supplementary Table S1). Those networks are composed of a disease monoplex network, a gene multiplex network and two genomic monoplex networks. The gene multiplex network encode gene interactions, and is identical to the one used in node prioritisation for leukemia. The disease monoplex network encodes disease phenotypic proximities. The two remaining networks encode genomic information: PCHi-C fragment interactions and TAD interactions. Importantly, the gene and disease networks are identical in the eight hematopoietic networks, whereas the PCHi-C and TAD networks were computed for each hematopoietic cell type from a dataset obtained in [20]. We describe in detail the processing pipeline applied to obtain the PCHi-C fragment and TAD layers in Supplementary Section 1.C.

### 4.2 RWR with MultiXrank

All RWR scores were obtained using the MultiXrank Python package [17], available on GitHub: https://github.com/anthbapt/multixrank. Parameters used in each study are described in Supplementary Section 2, 3 and 4 (Supplementary Tables S3, S6, S8 and S11).

### 4.3 Network visualisation

The network visualisation displayed in Figure 1 was obtained with Cytoscape [61].

### 4.4 Supervised classification

We created the gene-disease associations dataset from an outdated version of DisGeNET (v2.0, 2014). We obtained 1914 gene-disease associations. We generated a negative dataset by randomly picking 3828 pairs of gene and disease nodes that are not considered associated in DisGeNET v2.0 (2014). For each positive and negative gene-disease association defined in the training dataset, we used both the gene and the disease nodes as seeds when running MultiXrank (parameters defined in Supplementary Table S8). Then, we trained Random Forest and XGBoost binary classifiers (Supplementary Table S9) based on the MultiXrank output scores for predicting gene-disease associations. The evaluation of the binary classifiers is done on an updated version of the gene-disease associations dataset; DisGeNET v7.0 (2020). The test dataset contained 7218 novel positive gene-disease associations, and the negative dataset is randomly selected twice as many (i.e. 14 436) negative associations. We ran MultiXrank using as seeds the gene and disease nodes of each association of the evaluation dataset, using the same parameters used for obtaining MultiXrank scores for the training dataset (Supplementary Table S8). Then, the MultiXrank output scores were used as input of the previously computed Random Forest and XGBoost models (trained on the data obtained with the gene-disease association of DisGeNET v2.0 (2014)) to predict their label. Finally, we compared the predicted labels to the true labels. We report the results for each model in Supplementary Table S10. The full procedure is detailed in Supplementary Section 3 and Supplementary Figure S5.

### 4.5 Integration of MultiXrank output scores

#### Disease-disease rank distances per cell type and per node type

To compute disease-disease distance within each hematopoietic multilayer network, we start by integrating the scores of MultiXrank for each node type independently.

First, we create a disease distance matrix for each hematopoietic multilayer network *c* and node type *t* using equation 1:

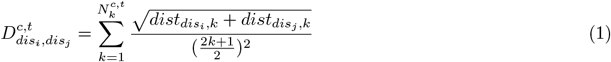

with *dis*_*i*_ and *dis*_*j*_, the two seeds (i.e. immune diseases) considered and 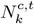 the number of nodes of type *t* in the multilayer network *c*.

The rank distances, 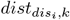 and 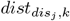, are computed using the following equations:

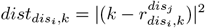

with 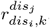, the rank of the node at position *k* for disease *dis*_*i*_ in the list of scores for disease *dis*_*j*_.

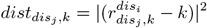

with 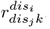, the rank of the node at position *k* for disease *dis*_*j*_ in the list of scores for disease *dis*_*i*_.

The distance 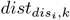 (respectively 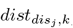) represents the square of the absolute difference between the rank of the node at position *k* for disease *dis*_*i*_ (resp. *dis*_*j*_) and the rank of the same node for disease *dis*_*j*_ (resp. *dis*_*i*_).

#### Disease-disease integrated rank distances

We integrate disease-disease rank distances across hematopoietic multilayer networks for each node type using the following equation:

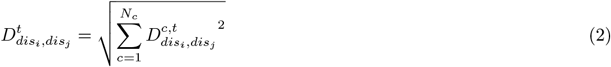

with 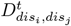 the integrated disease distances across hematopoietic multilayer networks for node type *t, N*_*c*_ the total number of hematopoietic multilayer networks (i.e. eight) and 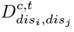 the disease-disease distances computed for hematopoietic multilayer network *c* and node type *t*.

### 4.6 Multiview Clustering

For clustering immune diseases based on the distance matrices 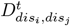 obtained for each node type *t*, we employed the multiview spectral clustering algorithm [62] implemented in the mvlearn python package [63] with *n clusters* = 3.

### 4.7 t-SNE projection

To visualise the immune disease clustering, we first concatenated the four immune disease distance matrices 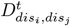. This concatenated matrix was then projected in a two-dimensional t-SNE (t-distributed stochastic neighbor embedding) space [40]. The points were colored according to their assigned cluster, obtained from the multiview clustering.

## Availability of data and materials

The MultiXrank package is available on GitHub: https://github.com/anthbapt/MultiXrank. MultiXrank can be installed with standard pip installation command: https://pypi.org/project/MultiXrank. It is associated with complete documentation: https://multixrank-doc.readthedocs.io/en/latest.

The code and data used in this study are available on Github: https://github.com/galadrielbriere/ApplicationsMultiXrank.

## Ethics approval and consent to participate

Not applicable.

## Consent for publication

Not applicable.

## Authors’ contributions

A.Bap. and A.Bau. designed research; A.Bap. an G.B performed research; A.Bap. and G.B analysed data; A.Bap. created numerical code, with contributions from G.B; A.Bap., G.B and A.Bau. wrote the paper.

## Additional Files

Additional file 1 — Supplementary information for: Random Walk with Restart on multilayer networks: from node prioritisation to supervised link prediction and beyond

Supplementary Section 1: Multilayer networks

Supplementary Section 2: Node prioritisation to study human genetic diseases Supplementary Section 3: Supervised prediction of gene-disease associations

Supplementary Section 4: Diffusion profiles comparison to unveil immune diseases similarities

Supplementary Table S1: Monoplex, multiplex and bipartite networks used in our three study cases. Of note, this table includes only the networks used in our analysis and does not encompass the broader non-network format information used in our studies.

Supplementary Table S2: PCHi-C fragment networks characteristics.

Supplementary Table S3: MultiXrank parameters used for node prioritisation in leukemia.

Supplementary Table S4: Top 10 prioritised genes for leukemia, associated MultiXrank scores, degree and distance to seed nodes.

Supplementary Table S5: Top 10 prioritised drugs for leukemia, associated MultiXrank scores, degree and distance to seed nodes.

Supplementary Table S6: Four sets of MultiXrank parameters used for node prioritisation in epilepsy. It is to note that only the *λ* parameter differs.

Supplementary Table S7: DrugBank Categories associated to at least 7 drugs from the 41 drugs prioritised by MultiXrank (parameter set number 4, top-100) that are not prioritised by Hetionet. The first column of the table corresponds to the DrugBank Category and the second column (N) corresponds to the number of drugs mapped to each category. It is to note that some drugs belong to more than one class.

Supplementary Table S8: MultiXrank parameters used for the supervised prediction of gene-disease associations.

Supplementary Table S9: XGBoost and Random Forest models and performance metrics for predicting Gene-Disease associations from DisGeNET v2.0 (2014). The models are trained and tested according to the protocol described in Supplementary Figure S5. We trained the XGBoost and Random Forest models using various class weights, large weights penalising classification errors for the corresponding class (pos., for positive associations and neg., for negative associations). We report several performance metrics for each model: Balanced Accuracy (defined as the average of recall obtained on each class), F1-score and number of True Positives (TP), False Positives (FP), True Negatives (TN) and False Negative (FN). The best performing model is highlighted in grey.

Supplementary Table S10: XGBoost and Random Forest models and performance metrics for predicting Gene-Disease associations from DisGeNET v7.0 (2020). The models are the same as those presented in

Supplementary Table S9, and have not been retrained for predicting DisGeNET v7.0 (2020) associations. We report several performance metrics for each model: Balanced Accuracy (defined as the average of recall obtained on each class), F1-score and number of True Positives (TP), False Positives (FP), True Negatives (TN) and False Negative (FN). The best performing model is highlighted in grey.

Supplementary Table S11: MultiXrank parameters used for diffusion profiles comparison to unveil immune diseases similarities.

Supplementary Table S12: Composition and characteristics of the immune disease clusters. Supplementary Table S13: List of the 131 immune diseases considered in this study. The first column represents the number used to identify each disease in the t-SNE projection. The second column represents the name of the disease. The third column is the UMLS identifier of the disease.

Supplementary Figure S1: Tree lineages of hematopoietic cells. MPP: Multi-Potent Progenitor ; CLP: Common Lymphoid Progenitor ; CMP: Common Myeloid Progenitor ; GMP: Granulocyte/Macrophage Progenitor ; MEP: Megakaryocyte/Erythrocyte Progenitor. Lymphoid cells and myeloid cells included in this study are indicated in red and blue, respectively.

Supplementary Figure S2: Left: Largest component of the PCHi-C fragment network built from the nB cells dataset. Right: Degree distribution of the PCHi-C fragment network defined on the left, with a zoom on low degree nodes. Supplementary Figure S3: Representation of the multilayer network used for node prioritisation in leukemia. The network is composed of a gene multiplex network, a drug multiplex network and their associated bipartite network. The gene multiplex network contains nodes corresponding to genes/proteins and edges corresponding to protein-protein, molecular complex, and Reactome pathway associations. The drug multiplex network contains nodes corresponding to drugs and edges corresponding to pharmacological, experimental, predicted, and clinical drug-drug interactions (Supplementary section 1.A). For the sake of clarity, the bipartite network containing the interactions connecting the two different types of nodes of the two multiplex networks is represented by the black arrow.

Supplementary Figure S4: Ictogenic properties of the 100 drugs prioritised by Hetionet and overlap with the top 100, 200, and 500 drugs prioritised by MultiXrank (parameter set 1 to 3, see Supplementary Table S6). The data on ictogenic properties has been sourced from the Hetionet study. AIGD are anti-ictogenic drugs that have a seizure suppressor effect (beige), IGD are ictogenic drugs (pink), and UKND are drugs with unknown effects (grey). The outer circle of the pie chart displays the number of AIGD, IGD, and UNKD drugs amongst the 100 drugs prioritised by Hetionet. The center of the pie chart displays the number of drugs from each category that were also prioritised by MultiXrank in the top-100 (displayed in lighter shades), top-200 (displayed in middle shade), and top-500 drugs (displayed in darker shades). The drugs prioritised in the top-500 include the drugs prioritised in the top-200, and the top-200 includes the drugs prioritised in the top-100. The white part of the pie chart corresponds to the drugs prioritised by Hetionet that are not in our MultiXrank top-500 prioritised drugs.

Supplementary Figure S5: Workflow of the random forest binary classifier. The left panel represents the training step. Here, the bipartite network connecting the gene multiplex and the disease monoplex network is built from DisGeNET v2.0 (2014). The gene-disease associations in DisGeNET v2.0 (2014) are also considered positive gene-disease associations. Negative gene-disease associations are sampled randomly. Then, MultiXrank is run using the gene and the disease nodes of each positive and negative gene-disease association, and the output scores are saved as described in the matrix *. This matrix is used with the positive and negative labels to train a random forest binary classifier. The right panel represents the test step. In this case, the true positive gene-disease associations are created from DisGeNET v7.0 (2020) and true negative gene-disease associations are sampled randomly. We next ran MultiXrank using as seeds the gene and disease nodes from each positive and negative gene-disease association, and saved the output scores in the matrix **. Finally, this ** matrix is used as an input of the previously trained random forest classifier to predict the labels of each gene-disease association. The predicted labels are then compared with the true known labels. The protocol is the same for two or three multiplex networks.

Supplementary Figure S6: 2D PCA projection of the Jaccard index similarities between the different hematopoietic cell types. Left: similarities computed on the PCHi-C fragment dataset. Right: similarities computed on the TAD dataset. Red: Lymphoid cells. Blue: Myeloid cells.

Supplementary Figure S7: 2D PCA projection of the similarity between the hematopoietic cell types. The tree lineage of hematopoietic cells is correctly found with the PCHi-C fragment, TAD and disease output scores of MultiXrank. However, the tree lineage is not recovered for the protein output scores.

Supplementary Figure S8: 2D PCA projection of hematopoietic cell similarities with respect to the integrated MultiXrank output scores effectively visualises the similarity between various hematopoietic cell types. In this projection, lymphoid cells (depicted in red) and myeloid cells (depicted in blue) exhibit a relative separation within the PCA space. Moreover, nCD4 and nCD8 cells are in close proximity to each other and relative proximity with nB cells. Furthermore, Macrophage (Mac0) and its precursor, Monocyte (Mon), appear closely situated. Additionally, Erythrocyte (Ery) and Megakaryocyte (MK), both stemming from the same progenitor, also exhibit high proximity to each other.

## References

1. Masuda, N., Porter, M.A., Lambiotte, R.: Random walks and diffusion on networks. Physics Reports 716-717, 1–58 (2017). doi:10.1016/j.physrep.2017.07.007. Random walks and diffusion on networks

2. Costa, L.d.F., Travieso, G.: Exploring complex networks through random walks. Phys. Rev. E 75, 016102 (2007). doi:10.1103/PhysRevE.75.016102

3. Macropol, K., Can, T., Singh, A.K.: Rrw: repeated random walks on genome-scale protein networks for local cluster discovery. BMC Bioinformatics 10(1), 283 (2009). doi:10.1186/1471-2105-10-283

4. Newman, M.E.J.: A measure of betweenness centrality based on random walks. Social Networks 27(1), 39–54 (2005). doi:10.1016/j.socnet.2004.11.009

5. Brin, S., Page, L.: The anatomy of a large-scale hypertextual web search engine. Computer Networks and ISDN Systems 30(1), 107–117 (1998). doi:10.1016/S0169-7552(98)00110-X. Proceedings of the Seventh International World Wide Web Conference

6. Pan, J.-Y., Yang, H.-J., Faloutsos, C., Duygulu, P.: Automatic multimedia cross-modal correlation discovery. In: Proceedings of the Tenth ACM SIGKDD International Conference on Knowledge Discovery and Data Mining. KDD ‘04, pp. 653–658. Association for Computing Machinery, New York, NY, USA (2004). doi:10.1145/1014052.1014135. https://doi.org/10.1145/1014052.1014135

7. Langville, A.N., Meyer, C.D.: Google’s PageRank and Beyond: The Science of Search Engine Rankings. Princeton University Press, USA (2006)

8. Gómez, S., Díaz-Guilera, A., Gómez-Gardeñes, J., Pérez-Vicente, C.J., Moreno, Y., Arenas, A.: Diffusion dynamics on multiplex networks. Phys. Rev. Lett. 110, 028701 (2013). doi:10.1103/PhysRevLett.110.028701

9. Köhler, S., Bauer, S., Horn, D., Robinson, P.N.: Walking the interactome for prioritization of candidate disease genes. The American Journal of Human Genetics 82(4), 949–958 (2008). doi:10.1016/j.ajhg.2008.02.013

10. Cho, H., Berger, B., Peng, J.: Compact integration of multi-network topology for functional analysis of genes. Cell Systems 3(6), 540–5485 (2016). doi:10.1016/j.cels.2016.10.017

11. Ko, Y., Cho, M., Lee, J.-S., Kim, J.: Identification of disease comorbidity through hidden molecular mechanisms. Scientific Reports 6(1), 39433 (2016). doi:10.1038/srep39433

12. Chen, X., Liu, M.-X., Yan, G.-Y.: Drug–target interaction prediction by random walk on the heterogeneous network. Mol. BioSyst. 8, 1970–1978 (2012). doi:10.1039/C2MB00002D

13. Peng, L., Shen, L., Xu, J., Tian, X., Liu, F., Wang, J., Tian, G., Yang, J., Zhou, L.: Prioritizing antiviral drugs against sars-cov-2 by integrating viral complete genome sequences and drug chemical structures. Scientific Reports 11(1), 6248 (2021). doi:10.1038/s41598-021-83737-5

14. Han, N., Hwang, W., Tzelepis, K., Schmerer, P., Yankova, E., MacMahon, M., Lei, W., Katritsis, N.M., Liu, A., Felgenhauer, U., Schuldt, A., Harris, R., Chapman, K., McCaughan, F., Weber, F., Kouzarides, T.: Identification of sars-cov-2-induced pathways reveals drug repurposing strategies. Science Advances 7(27), 3032 (2021). doi:10.1126/sciadv.abh3032. https://www.science.org/doi/pdf/10.1126/sciadv.abh3032

15. Li, Y., Patra, J.C.: Genome-wide inferring gene–phenotype relationship by walking on the heterogeneous network. Bioinformatics 26(9), 1219–1224 (2010). doi:10.1093/bioinformatics/btq108. https://academic.oup.com/bioinformatics/article-pdf/26/9/1219/29012989/btq108.pdf

16. Valdeolivas, A., Tichit, L., Navarro, C., Perrin, S., Odelin, G., Levy, N., Cau, P., Remy, E., Baudot, A.: Random walk with restart on multiplex and heterogeneous biological networks. Bioinformatics 35(3), 497–505 (2018). doi:10.1093/bioinformatics/bty637. https://academic.oup.com/bioinformatics/article-pdf/35/3/497/27699899/bty637.pdf

17. Baptista, A., Gonzalez, A., Baudot, A.: Universal multilayer network exploration by random walk with restart. Communications Physics 5(1), 170 (2022). doi:10.1038/s42005-022-00937-9

18. Himmelstein, D.S., Lizee, A., Hessler, C., Brueggeman, L., Chen, S.L., Hadley, D., Green, A., Khankhanian, P., Baranzini, S.E.: Systematic integration of biomedical knowledge prioritizes drugs for repurposing. eLife 6, 26726 (2017). doi:10.7554/eLife.26726

19. Schoenfelder, S., Furlan-Magaril, M., Mifsud, B., Tavares-Cadete, F., Sugar, R., Javierre, B.-M., Nagano, T., Katsman, Y., Sakthidevi, M., Wingett, S.W., Dimitrova, E., Dimond, A., Edelman, L.B., Elderkin, S., Tabbada, K., Darbo, E., Andrews, S., Herman, B., Higgs, A., LeProust, E., Osborne, C.S., Mitchell, J.A., Luscombe, N.M., Fraser, P.: The pluripotent regulatory circuitry connecting promoters to their long-range interacting elements. Genome research 25, 582–97 (2015)

20. Javierre, B.M., Burren, O.S., Wilder, S.P., Kreuzhuber, R., Hill, S.M., Sewitz, S., Cairns, J., Wingett, S.W., Várnai, C., Thiecke, M.J., Burden, F., Farrow, S., Cutler, A.J., Rehnström, K., Downes, K., Grassi, L., Kostadima, M., Freire-Pritchett, P., Wang, F., Martens, J.H., Kim, B., Sharifi, N., Janssen-Megens, E.M., Yaspo, M.-L., Linser, M., Kovacsovics, A., Clarke, L., Richardson, D., Datta, A., Flicek, P., Stunnenberg, H.G., Todd, J.A., Zerbino, D.R., Stegle, O., Ouwehand, W.H., Frontini, M., Wallace, C., Spivakov, M., Fraser, P.: Lineage-specific genome architecture links enhancers and non-coding disease variants to target gene promoters. Cell 167(5), 1369–138419 (2016). doi:10.1016/j.cell.2016.09.037

21. Tyner, J.W., Erickson, H., Deininger, M.W.N., Willis, S.G., Eide, C.A., Levine, R.L., Heinrich, M.C., Gattermann, N., Gilliland, D.G., Druker, B.J., Loriaux, M.M.: High-throughput sequencing screen reveals novel, transforming ras mutations in myeloid leukemia patients. Blood 113(19075190), 1749–1755 (2009)

22. Thomas, X., Elhamri, M.: Tipifarnib in the treatment of acute myeloid leukemia. Biologics : targets & therapy 1(19707311), 415–424 (2007)

23. Yanamandra, N., Buzzeo, R.W., Gabriel, M., Hazlehurst, L.A., Mari, Y., Beaupre, D.M., Cuevas, J.: Tipifarnib-induced apoptosis in acute myeloid leukemia and multiple myeloma cells depends on ca2+ influx through plasma membrane ca2+ channels. J Pharmacol Exp Ther 337(3), 636 (2011)

24. Luger, S., Wang, V.X., Paietta, E., Ketterling, R.P., Rybka, W., Lazarus, H.M., Litzow, M.R., Rowe, J.M., Larson, R.A., Appelbaum, F.R., Tallman, M.S.: Tipifarnib as maintenance therapy in acute myeloid leukemia (aml) improves survival in a subgroup of patients with high risk disease. results of the phase iii intergroup trial e2902. Blood 126(23), 1308–1308 (2015). doi:10.1182/blood.V126.23.1308.1308

25. McGeady, P., Kuroda, S., Shimizu, K., Takai, Y., Gelb, M.H.: The farnesyl group of h-ras facilitates the activation of a soluble upstream activator of mitogen-activated protein kinase. The Journal of biological chemistry 270, 26347–51 (1995)

26. Su, M., Chang, Y.-T., Hernandez, D., Jones, R.J., Ghiaur, G.: Regulation of drug metabolizing enzymes in the leukaemic bone marrow microenvironment. Journal of Cellular and Molecular Medicine 23(6), 4111–4117 (2019). doi:10.1111/jcmm.14298. eprint: https://onlinelibrary.wiley.com/doi/pdf/10.1111/jcmm.14298. Accessed 2023-07-04

27. Venkatasubbarao, K., Choudary, A., Freeman, J.W.: Farnesyl transferase inhibitor (R115777)-induced inhibition of STAT3(Tyr705) phosphorylation in human pancreatic cancer cell lines require extracellular signal-regulated kinases. Cancer Research 65(7), 2861–2871 (2005). doi:10.1158/0008-5472.CAN-04-2396

28. Laverdiere, I., Boileau, M., Neumann, A., Frison, H., Mitchell, A., Ng, S., Wang, J., Minden, M., Eppert, K.: Leukemic stem cell signatures identify novel therapeutics targeting acute myeloid leukemia. Blood Cancer Journal 8 (2018). doi:10.1038/s41408-018-0087-2

29. Matsumoto, S., Yamazoe, Y.: Involvement of multiple human cytochromes P450 in the liver microsomal metabolism of astemizole and a comparison with terfenadine. British Journal of Clinical Pharmacology 51(2), 133–142 (2001). doi:10.1111/j.1365-2125.2001.01292.x. Accessed 2023-07-04

30. Wishart, D.S., Feunang, Y.D., Guo, A.C., Lo, E.J., Marcu, A., Grant, J.R., Sajed, T., Johnson, D., Li, C., Sayeeda, Z., Assempour, N., Iynkkaran, I., Liu, Y., Maciejewski, A., Gale, N., Wilson, A., Chin, L., Cummings, R., Le, D., Pon, A., Knox, C., Wilson, M.: DrugBank 5.0: a major update to the DrugBank database for 2018. Nucleic Acids Research 46(D1), 1074–1082 (2018). doi:10.1093/nar/gkx1037

31. Runtz, L., Girard, B., Toussenot, M., Espallergues, J., Fayd’Herbe De Maudave, A., Milman, A., deBock, F., Ghosh, C., Guérineau, N.C., Pascussi, J.-M., Bertaso, F., Marchi, N.: Hepatic and hippocampal cytochrome p450 enzyme overexpression during spontaneous recurrent seizures. Epilepsia 59, 123–134 (2018)

32. Gogou, M., Pavlou, E.: Efficacy of antiepileptic drugs in the era of pharmacogenomics: A focus on childhood. European Journal of Paediatric Neurology 23(5), 674–684 (2019). doi:10.1016/j.ejpn.2019.06.004. Accessed 2023-07-05

33. Wilner, A.N., Sharma, B.K., Soucy, A., Thompson, A., Krueger, A.: Common comorbidities in women and men with epilepsy and the relationship between number of comorbidities and health plan paid costs in 2010. Epilepsy & Behavior: E&B 32, 15–20 (2014). doi:10.1016/j.yebeh.2013.12.032

34. Stöllberger, C., Finsterer, J.: Cardiorespiratory findings in sudden unexplained/unexpected death in epilepsy (SUDEP). Epilepsy Research 59(1), 51–60 (2004). doi:10.1016/j.eplepsyres.2004.03.008

35. Szczurkowska, P.J., Polonis, K., Becari, C., Hoffmann, M., Narkiewicz, K., Chrostowska, M.: Epilepsy and hypertension: The possible link for sudden unexpected death in epilepsy? Cardiology Journal 28(2), 330–335 (2021). doi:10.5603/CJ.a2019.0095. Accessed 2023-07-05

36. Ata, S.K., Wu, M., Fang, Y., Ou-Yang, L., Kwoh, C.K., Li, X.-L.: Recent advances in network-based methods for disease gene prediction. Briefings in bioinformatics (2020). doi:10.1093/bib/bbaa303

37. Piñero, J., Queralt-Rosinach, N., Bravo, A., Deu-Pons, J., Bauer-Mehren, A., Baron, M., Sanz, F., Furlong, L.I.: Disgenet: a discovery platform for the dynamical exploration of human diseases and their genes. Database (Oxford) 2015 (2015)

38. Piñero, J., Ramírez-Anguita, J.M., Saüch-Pitarch, J., Ronzano, F., Centeno, E., Sanz, F., Furlong, L.I.: The disgenet knowledge platform for disease genomics: 2019 update. Nucleic Acids Res 48(D1), 845–855 (2020)

39. Spielmann, M., Lupiáñez, D.G., Mundlos, S.: Structural variation in the 3d genome. Nature Reviews Genetics 19(7), 453–467 (2018). doi:10.1038/s41576-018-0007-0

40. van der Maaten, L., Hinton, G.: Visualizing data using t-sne. Journal of Machine Learning Research 9(86), 2579–2605 (2008)

41. Hedrich, C.M., Tsokos, G.C.: Bridging the gap between autoinflammation and autoimmunity. Clinical Immunology 147(3), 151–154 (2013). doi:10.1016/j.clim.2013.03.006. Accessed 2023-09-27

42. Hedrich, C.M.: Shaping the spectrum — From autoinflammation to autoimmunity. Clinical Immunology 165, 21–28 (2016). doi:10.1016/j.clim.2016.03.002. Accessed 2023-09-27

43. Hsing, A.W., Hansson, L.-E., McLaughlin, J.K., Nyren, O., Blot, W.J., Ekbom, A., Fraumeni Jr., J.F.: Pernicious anemia and subsequent cancer. A population-based cohort study. Cancer 71(3), 745–750 (1993). doi:10.1002/1097-0142(19930201)71:3¡745::AID-CNCR2820710316¿3.0.CO;2-1. eprint: https://onlinelibrary.wiley.com/doi/pdf/10.1002/1097-0142%2819930201%2971%3A3%3C745%3A%3AAID-CNCR2820710316%3E3.0.CO%3B2-1. Accessed 2023-09-27

44. Corey, S.J., Minden, M.D., Barber, D.L., Kantarjian, H., Wang, J.C.Y., Schimmer, A.D.: Myelodysplastic syndromes: the complexity of stem-cell diseases. Nature Reviews Cancer 7(2), 118–129 (2007). doi:10.1038/nrc2047. Number: 2 Publisher: Nature Publishing Group. Accessed 2023-09-27

45. Taylor, A.M., Metcalfe, J.A., Thick, J., Mak, Y.F.: Leukemia and lymphoma in ataxia telangiectasia. Blood 87(2), 423–438 (1996)

46. Arora, H., Chacon, A.H., Choudhary, S., McLeod, M.P., Meshkov, L., Nouri, K., Izakovic, J.: Bloom syndrome. International Journal of Dermatology 53(7), 798–802 (2014). doi:10.1111/ijd.12408. eprint: https://onlinelibrary.wiley.com/doi/pdf/10.1111/ijd.12408. Accessed 2023-09-27

47. Mäkitie, O., Pukkala, E., Teppo, L., Kaitila, I.: Increased incidence of cancer in patients with cartilage-hair hypoplasia. The Journal of Pediatrics 134(3), 315–318 (1999). doi:10.1016/S0022-3476(99)70456-7. Accessed 2023-09-27

48. Argyle, J.C., Kjeldsberg, C.R., Marty, J., Shigeoka, A.O., Hill, H.R.: T-Cell Lymphoma and the Chediak-Higashi Syndrome. Blood 60(3), 672–676 (1982). doi:10.1182/blood.V60.3.672.672. Accessed 2023-09-27

49. Sanlaville, D., Verloes, A.: CHARGE syndrome: an update. European Journal of Human Genetics 15(4), 389–399 (2007). doi:10.1038/sj.ejhg.5201778. Number: 4 Publisher: Nature Publishing Group. Accessed 2023-09-27

50. Niceta, M., Barresi, S., Pantaleoni, F., Capolino, R., Dentici, M.L., Ciolfi, A., Pizzi, S., Bartuli, A., Dallapiccola, B., Tartaglia, M., Digilio, M.C.: TARP syndrome: Long-term survival, anatomic patterns of congenital heart defects, differential diagnosis and pathogenetic considerations. European Journal of Medical Genetics 62(6), 103534 (2019). doi:10.1016/j.ejmg.2018.09.001. Accessed 2023-09-27

51. Goldmuntz, E.: 22q11.2 deletion syndrome and congenital heart disease. American Journal of Medical Genetics. Part C, Seminars in Medical Genetics 184(1), 64–72 (2020). doi:10.1002/ajmg.c.31774

52. Shah, S.S., Chhabra, M.: Parry-Romberg Syndrome. In: StatPearls. StatPearls Publishing, Treasure Island (FL) (2023). http://www.ncbi.nlm.nih.gov/books/NBK574506/ Accessed 2023-09-27

53. Ranum, A., Freese, R., Ramesh, V., Pearson, D.R.: Lichen sclerosus in female patients is associated with an increased risk of metabolic syndrome and cardiovascular comorbidities: a retrospective cohort review. British Journal of Dermatology 187(6), 1030–1032 (2022). doi:10.1111/bjd.21811. Accessed 2023-09-27

54. Shahid, S., El Assaad, I., Patel, A., Parikh, S., Aziz, P.F.: Conduction defects in pediatric patients with Pearson syndrome: When to pace? Heart Rhythm, 1547–527123024141 (2023). doi:10.1016/j.hrthm.2023.07.004

55. Ballo, P., Chiodi, L., Cameli, M., Malandrini, A., Federico, A., Mondillo, S., Zuppiroli, A.: Dilated cardiomyopathy and inclusion body myositis. Neurological Sciences: Official Journal of the Italian Neurological Society and of the Italian Society of Clinical Neurophysiology 33(2), 367–370 (2012). doi:10.1007/s10072-011-0766-2

56. Chan, Y.C., Lee, Y.S., Wong, S.T., Lam, S.P., Ong, B.K.C., Wilder-Smith, E.: Melkerrson–Rosenthal syndrome with cardiac involvement. Journal of Clinical Neuroscience 11(3), 309–311 (2004). doi:10.1016/j.jocn.2003.06.003. Accessed 2023-09-27

57. Ferguson, P.J., El-Shanti, H.: Majeed Syndrome: A Review of the Clinical, Genetic and Immunologic Features. Biomolecules 11(3), 367 (2021). doi:10.3390/biom11030367. Number: 3 Publisher: Multidisciplinary Digital Publishing Institute. Accessed 2023-09-28

58. Bellon, N., Paluel-Marmont, C., De Peufeilhoux, L., Barbet, P., Bodemer, C., Dupont, C.: Eosinophilic esophagitis is a trait of netherton syndrome. Journal of Allergy and Clinical Immunology 137(2), 280 (2016). doi:10.1016/j.jaci.2015.12.1165

59. Paik, J.J., Corse, A.M., Mammen, A.L.: The Co-Existence of Myasthenia Gravis in Patients with Myositis: A Case Series. Seminars in arthritis and rheumatism 43(6), 792–796 (2014). doi:10.1016/j.semarthrit.2013.12.005. Accessed 2023-09-28

60. Brown, A.S., Patel, C.J.: A standard database for drug repositioning. Scientific Data 4(1), 170029 (2017). doi:10.1038/sdata.2017.29

61. Shannon, P., Markiel, A., Ozier, O., Baliga, N.S., Wang, J.T., Ramage, D., Amin, N., Schwikowski, B., Ideker, T.: Cytoscape: a software environment for integrated models of biomolecular interaction networks. Genome Research 13(11), 2498–2504 (2003). doi:10.1101/gr.1239303

62. Kumar, A., III, H.D.: A co-training approach for multi-view spectral clustering. In: Proceedings of the 28th International Conference on International Conference on Machine Learning. ICML’11, pp. 393–400. Omnipress, Madison, WI, USA (2011)

63. Perry, R., Mischler, G., Guo, R., Lee, T., Chang, A., Koul, A., Franz, C., Richard, H., Carmichael, I., Ablin, P., Gramfort, A., Vogelstein, J.T.: mvlearn: Multiview machine learning in python. Journal of Machine Learning Research 22(109), 1–7 (2021)

