## Supplementary materials for "Random Walk with Restart on multilayer networks: from node prioritisation to supervised link prediction and beyond"

##### This PDF file includes:

Figs. S1 to S8
Tables S1 to S13
SI References

### 1. Multilayer networks

#### A. Biological multilayer network.

1. Gene multiplex network. This multiplex network is composed of three layers of interactions between nodes corresponding to genes or proteins, here considered indifferently, and named according to their official gene names.
   - Complexes: A molecular complexes layer constructed from the fusion of Hu.map (2) and Corum (3), using OmniPathR (4).
   - PPI: A Protein-Protein interaction (PPI) layer corresponding to the fusion of 3 datasets: APID (homo sapiens level 2, without inter-species interactions), Hi-Union and Lit-BM (www.interactome-atlas.org/download). For APID proteins that did not match any official gene names, we used their corresponding Uniprot IDs, which was the case for 166 over 13 346 APID proteins.
   - Reactome: a pathways layer extracted from NDEx (5) and corresponding to human Reactome data (6). The network is available in NDEx: https://www.ndexbio.org/viewer/networks/4bc71515-86da-11e7-a10d-0ac135e8bacf.
2. Disease monoplex network. In this network, the nodes correspond to diseases, labelled by their UMLS identifiers, and the edges were built based on phenotypic proximities between diseases, computed considering the set of their associated phenotypes in the Human Phenotype Ontology (7). The protocol for computing those phenotypic proximities is detailed in (8).
3. Drug multiplex network. This multiplex network is composed of four layers of interactions between nodes representing drugs, identified with their DrugBank (9) names. The interaction data of this multiplex network were taken from Cheng et al. (10)† (first three layers) and from https://snap.stanford.edu/biodata/datasets/10001/10001-ChCh-Miner.html‡ (fourth layer).
   - Drug1†: Clinical drug interactions (clinically reported adverse drug-drug interactions, 14822 clinically reported adverse drug–drug interactions connecting 667 drugs.)
   - Drug2†: Experimental drug combinations (experimentally validated drug combinations, 737 unique pairwise drug combinations connecting 376 drugs.)
   - Drug3†: Predicted drug combinations (network-predicted hypertensive drug combinations, 2080 potential combinations involving 65 hypertensive drugs.)
   - Drug4‡: Drug-Drug interactions determined from the pharmacological effect of the action of one drug on another drug (48514 interactions involving 1514 drugs).
4. Promoter Capture Hi-C (PCHi-C) fragment monoplex networks. These networks represent the interactions between sequences of DNA. The nodes represents DNA fragments, i.e. a start and an end position on a chromosome sequence. Two fragments nodes are connected with an edge if the two DNA sequences interacts. These networks were build from the dataset provided by Javierre and co-authos (11). Importantly, we generated 8 PCHi-C fragment networks, one for each of the hematopoietic cell lines considered in our studiy. However, those 8 networks are not used in the same multilayer network, hence they are considered as monoplex networks. We detail the construction of these hematopoietic cell line specific PCHi-C fragment networks in Supplementary Section 1.C.
5. Topologically Associated Domains (TAD) monoplex networks. Thse networks represent the adjacency of TADs. The nodes represent TADs, defined by a start and end position of a chromosome sequence, are edges connects adjacent TADs. These networks were build from the dataset provided by Javierre and co-authos (11). Importantly, we generated 8 TAD networks, one for each of the hematopoietic cell lines considered in our study. However, those 8 networks are not used in the same multilayer network, hence they are considered as monoplex networks. We detail the construction of these hematopoietic cell line specific TAD networks in Supplementary Section 1.C.

**Table S1.** Monoplex, multiplex and bipartite networks used in our three study cases. Of note, this table includes only the networks used in our analysis and does not encompass the broader non-network format information used in our studies.

<table>
<tr><th rowspan="2">Type</th><th rowspan="2">Name</th><th rowspan="2">Association</th><th colspan="2">Study 1: Node prioritisation</th><th rowspan="2">Study 2: Gene-Disease association prediction</th><th rowspan="2">Study 3: Comparison of diseases diffusion profiles</th></tr>
<tr><th>Leukemia</th><th>Epilepsy</th></tr>
<tr><td rowspan="11">Monoplex and multiplex networks</td><td rowspan="3">Gene multiplex</td><td>Molecular complexes membership</td><td rowspan="3">X</td><td rowspan="3"></td><td rowspan="3">X</td><td rowspan="3">X</td></tr>
<tr><td>Protein-Protein interactions</td></tr>
<tr><td>Pathway membership</td></tr>
<tr><td>Disease monoplex</td><td>Phenotypic proximity</td><td></td><td></td><td>X</td><td>X</td></tr>
<tr><td rowspan="4">Drug multiplex</td><td>Clinically reported drug-drug interactions (adverse events)</td><td rowspan="4">X</td><td rowspan="4"></td><td rowspan="4"></td><td rowspan="4"></td></tr>
<tr><td>Experimental drug-drug combinations</td></tr>
<tr><td>Predicted drug-drug combinations</td></tr>
<tr><td>Drug-drug pharmacological interactions</td></tr>
<tr><td>PCHi-C fragment monoplex</td><td>Interactions between chromatin segments for 8 hematopoietic cell types (one network per cell type)</td><td></td><td></td><td></td><td>X</td></tr>
<tr><td>TAD monoplex</td><td>TAD adjacency in the genome for 8 hematopoeietic cell types (one network per cell type)</td><td></td><td></td><td></td><td>X</td></tr>
<tr style="display:none"></tr>
<tr><td rowspan="6">Bipartite networks</td><td>Gene-Disease DisGeNET v2.0 (2014)</td><td rowspan="2">Curated, inferred and literature extracted associations, scored according to their level of evidence</td><td></td><td></td><td>X</td><td></td></tr>
<tr><td>Gene-Disease DisGeNET v7.0 (2020)</td><td></td><td></td><td></td><td>X</td></tr>
<tr><td>Gene-Drug</td><td>Drug-Target associations</td><td>X</td><td></td><td></td><td></td></tr>
<tr><td>Gene-Fragment</td><td>Genes associations to promoters in the fragment</td><td></td><td></td><td></td><td>X</td></tr>
<tr><td>Gene-TAD</td><td>Gene included in TAD region</td><td></td><td></td><td></td><td>X</td></tr>
<tr><td>Fragment-TAD</td><td>Fragment included in TAD region</td><td></td><td></td><td></td><td>X</td></tr>
<tr><td colspan="3">Hetionet network (v1.0) (1)</td><td></td><td>X</td><td></td><td></td></tr>
</table>

These various multiplex and monoplex networks can be connected to each other with bipartite networks, defined as follows:

- Gene-Disease (1-2): Gene-disease associations were downloaded from the DisGeNET database, which contains associations derived from various sources and scored according to their level of evidence (formula reported in https://www.DisGeNET.org/dbinfo). Two version of this bipartite network were used in our study:
  - Outdated: Associations derived from an outdated version of DisGeNET (v2.0, 2014) (12).
  - Updated: Associations derived from an updated version of DisGeNET (v7.0, 2020) (13).
- Gene-Drug (1-3): Drug-target associations were downloaded from several sources, and merged to obtain this bipartite network: DrugBank v5.1.8 go.drugbank.com/releases/latest, DrugCentral v10.12 drugcentral.org/download, and Cheng et al (10).
- Disease-Drug (2-3): Approved drug-disease indications were extracted from repoDB (14).
- Gene-PCHi-C Fragment (1-4): Gene-PCHi-C fragment associations were defined from the article of Javierre et al. (11), which reported genes associated to each PCHi-C fragment using the version GRCh37.p13 of the genome.
- Gene-TAD (1-5): This bipartite network establishes an association between a gene and a TAD when the gene is located within the boundaries of the TAD. Those associations were computed using gencode V19 (15), and the GRCh37.p13 version of the genome.
- PCHi-C fragment-TAD (4-5): This bipartite network connects TAD and PCHi-C fragments if the PCHi-C fragment is located within the boundaries of the TAD.

Because MultiXrank only scores nodes appearing in multiplex networks, nodes that did not share any interaction other than bipartite (i.e. with no interaction in any of the monoplex networks composing a multiplex network) were artificially added to their corresponding multiplex network by creating self-interactions (i.e. self-loop). This intervention is purely technical and does not have effects on MultiXrank output scores.

##### B. Hetionet multilayer network.

The Hetionet network is a gene-, compound- and disease-centric network constructed by integrating knowledge and experimental findings from millions of biomedical research publications (1).

Hetionet is composed of three layers:

- Gene multiplex network composed of three layers: Gene–covaries–Gene (GcG), Gene–interacts–Gene (GiG), Gene-regulates-Gene (GrG)
- Disease monoplex network: Disease–resembles–Disease (DrD)
- Compound monoplex network: Compound–resembles–Compound (CrC)

Gene, disease and compounds nodes are connected by several bipartite networks:

- Gene-Disease: Union of three types of interaction, Disease–associates–Gene (DaG), Disease–downregulates–Gene (DdG), Disease–upregulates–Gene (DuG)
- Compound-Disease: Union of two types of interaction, Compound–palliates–Disease (CpD), Compound–treats–Disease (CtD)
- Compound-Gene: Union of two types of interaction, Compound–downregulates–Gene (CdG), Compound–upregulates–Gene (CuG)

Hetionet also consider additional bipartite interactions for genes, compounds and diseases:

- Gene-Pathway: Gene–participates–Pathway (GpPW)
- Gene-Biological Process: Gene–participates–Biological Process (GpBP)
- Gene-Molecular Function: Gene–participates–Molecular Function (GpMF)
- Gene-Cellular Component: Gene–participates–Cellular Component (GpCC)
- Anatomy Part-Gene: Merged of three layers, Anatomy–downregulates–Gene (AdG), Anatomy–expresses–Gene (AeG), Anatomy–upregulates–Gene (AuG)
- Disease-Anatomy part: Disease–localises–Anatomy (DlA)
- Disease-Symptom: Disease–presents–Symptom (DpS)
- Compound-Side effect: Compound–causes–Side Effect (CcSE)
- Pharmalogic classes-Compound: Pharmacologic Class–includes–Compound (PCiC)

Since only bipartite interactions are considered for many node types (Pathway, Biological Process, Anatomy parts, Symptoms, Side Effects and Pharmacologic class), for these, we created artificial self-loops to match MultiXrank formalism. This intervention is purely technical and does not have effects on MultiXrank output scores.

#### C. Construction of the PCHi-C fragment and TAD networks.

The PCHi-C fragment and TAD networks were generated for various hematopoietic cell lines, based on the datasets generated by Javierre et al. (11). Eight hematopoietic cell lines were considered: Erythrocytes (Ery), Macrophages (Mac0), Megakaryocytes (MK), Monocytes (Mon), Naive B Cells (nB), Naive CD4+ T Cells (nCD4), Naive CD8+ T Cells (nCD8) and Neutrophils (Neu). The corresponding tree lineage is reported in Supplementary Figure S1.

The datasets, originally produced and processed by Javierre and co-authors using the CHICAGO pipeline (16) to identify significant interactions (score $> 5$) in various hematopoietic cell lines. The processed data are represented below:

| fragments 1 | | | fragments 2 | | | Score |
|---|---|---|---|---|---|---|
| chr | start | end | chr | start | end | |
| | ⋮ | | | ⋮ | | ⋮ |
| | ⋮ | | | ⋮ | | ⋮ |
| | ⋮ | | | ⋮ | | ⋮ |

Using these results, we were able to create a network for each hematopoietic cell line, containing the PCHi-C fragment interactions identified by Javierre et al.

In their study, Javierre et al. also used the directionality index score (17) to identify TADs in the 8 hematopoietic cell lines. The dataset is represented below:

| TADs | | |
|---|---|---|
| chr | start | end |
| | ⋮ | |
| | ⋮ | |
| | ⋮ | |

Building upon these results, we were able to construct the TAD networks by connecting adjacent TADs.

We report several features of the obtained PCHi-C fragment networks in Supplementary Table S2 and in Supplementary Figure S2.

**Table S2.** PCHi-C fragment networks characteristics

| Cellular Type | Naive B cells (nB) | Erythroblasts (Ery) | Macrophages M0 (Mac0) | Monocytes (Mon) |
|---|---|---|---|---|
| # Nodes | 101973 | 90184 | 109566 | 96443 |
| # Edges | 192104 | 152617 | 182250 | 167275 |
| # Isolated Nodes | 0 | 0 | 0 | 0 |
| # Self Loops | 0 | 0 | 0 | 0 |
| Density | 3.7E-05 | 3.8E-05 | 3E-05 | 3.6E-05 |
| Avg. Clustering Coeff. | 0.152 | 0.092 | 0.094 | 0.14 |
| Avg. Degree | 3.768 | 3.385 | 3.327 | 3.469 |

| Cellular Type | Naive CD4+ T cells (nCD4) | Naive CD8+ T cells (nCD8) | Neutrophils (Neu) | Megakaryocytes (MK) |
|---|---|---|---|---|
| # Nodes | 106849 | 108016 | 81244 | 98159 |
| # Edges | 212106 | 218293 | 143679 | 152347 |
| # Isolated Nodes | 0 | 0 | 0 | 0 |
| # Self Loops | 0 | 0 | 0 | 0 |
| Density | 3.7E-05 | 3.7E-05 | 4.4E-05 | 3.2E-05 |
| Avg. Clustering Coeff. | 0.188 | 0.18 | 0.151 | 0.091 |
| Avg. Degree | 3.97 | 4.042 | 3.537 | 3.104 |

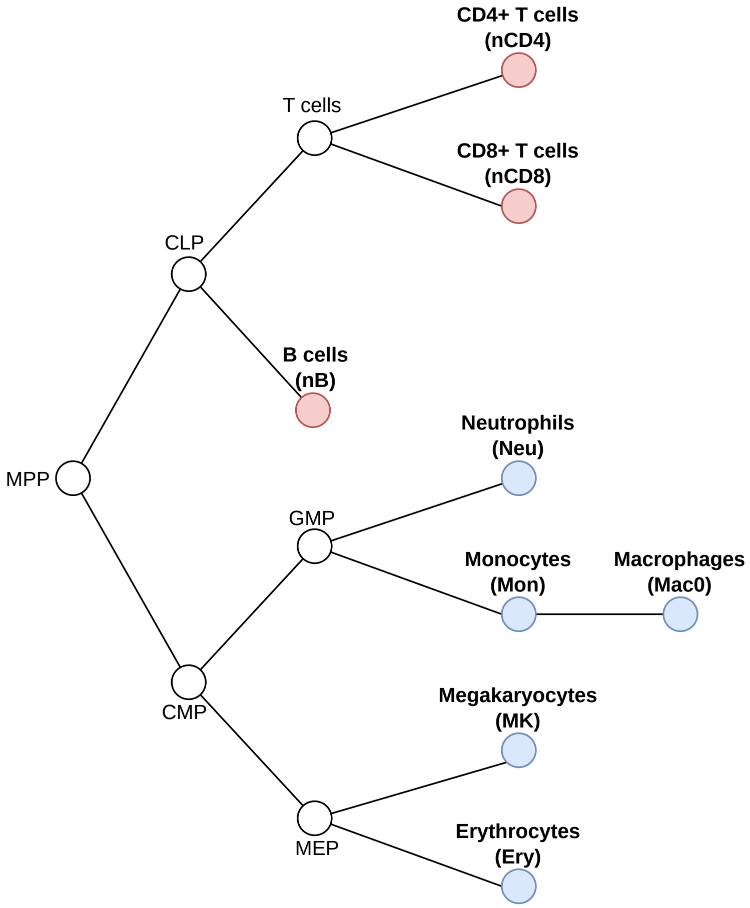

**Fig. S1.** Tree lineages of hematopoietic cells. MPP: Multi-Potent Progenitor ; CLP: Common Lymphoid Progenitor ; CMP: Common Myeloid Progenitor ; GMP: Granulocyte/Macrophage Progenitor ; MEP: Megakaryocyte/Erythrocyte Progenitor. Lymphoid cells and myeloid cells included in this study are indicated in red and blue, respectively.

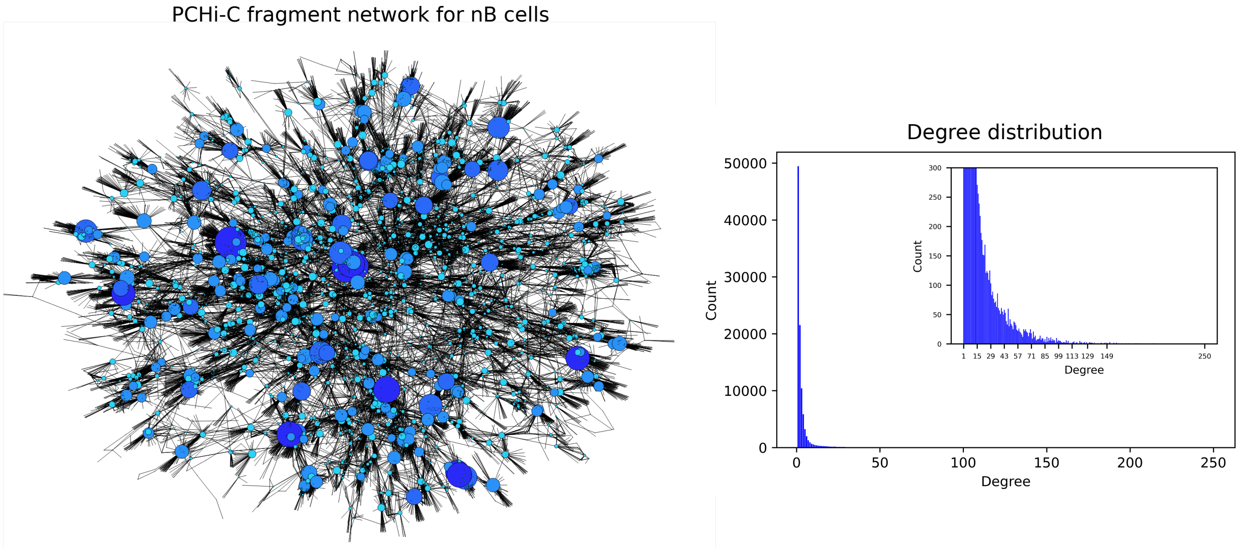

**Fig. S2.** Left: Largest component of the PCHi-C fragment network built from the nB cells dataset. Right: Degree distribution of the PCHi-C fragment network defined on the left, with a zoom on low degree nodes.

#### 2. Node prioritisation to study human genetic diseases

##### A. Node prioritisation in Leukemia.

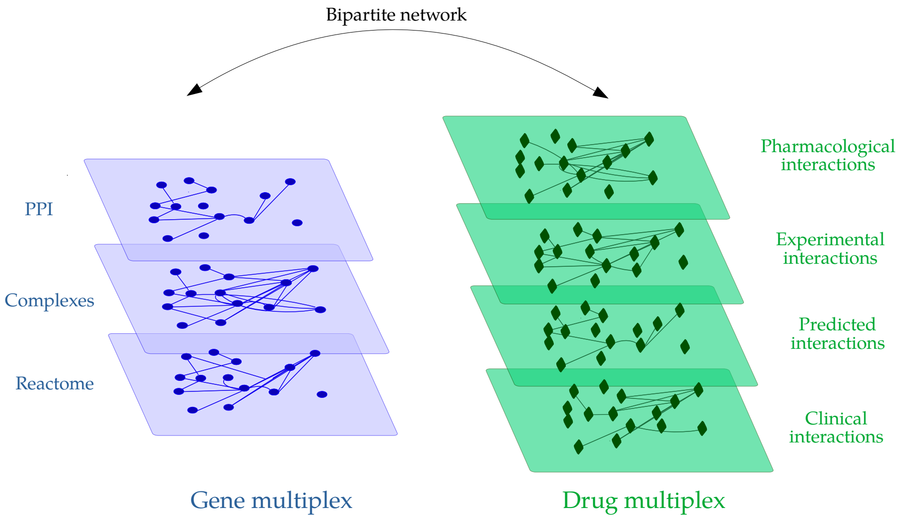

**Fig. S3.** Representation of the multilayer network used for node prioritisation in leukemia. The network is composed of a gene multiplex network, a drug multiplex network and their associated bipartite network. The gene multiplex network contains nodes corresponding to genes/proteins and edges corresponding to protein-protein, molecular complex, and Reactome pathway associations. The drug multiplex network contains nodes corresponding to drugs and edges corresponding to pharmacological, experimental, predicted, and clinical drug-drug interactions (Supplementary section 1.A). For the sake of clarity, the bipartite network containing the interactions connecting the two different types of nodes of the two multiplex networks is represented by the black arrow.

Top-10 prioritised genes and their known or suspected relations with leukemia:

1. CYP3A4 is known to be implicated in leukemia (18).
2. FNTB is the target of Tipifarnib (our seed drug) according to the DrugBank database.
3. RAF1 mutations were identified in patients with Noonan syndrome (19), which is a well-known risk factor for leukemia (20).
4. RASGRP1 has been linked with leukemia (21).
5. DGKZ knockdown can induce apoptosis in human Acute Myeloid Leukemia (AML) HL-60 cells through the MAPK/survivin/caspase pathway (22).
6. RIN1 plays a role in the maintenance of the abnormal RTK signaling in chronic myeloid leukemia (23).
7. BRAF mutations were identified in AML patients with monocytic differentiation (24).
8. RASA1 is involved in several types of cancer, including leukemia (25).
9. AURKA: A significant association between over-expression of AURKA and cytogenetic abnormalities was found in AML patients (26).
10. ARAF: A study found ARAF gene mutation in MOLT-4 leukemia cell line (27).

Top-10 prioritised drugs and their known or suspected relations with leukemia:

1. DB00637 (Astemizole) belongs to a drug class that was validated for its activities against human primary AML samples (28).

2. DB01380 (Cortisone acetate) is associated to acute leukemia, according to the DrugBank database.
3. DB00630 (Alendronate): Some preliminary experiments show a benefit for the treatment of osteopenia/osteoporosis with Alendronate in children with acute lymphoblastic leukemia (29).
4. DB00398 (Sorafenib) is a drug used for the treatment of advanced renal cell carcinoma, and some studies mention its impact on leukemia (30).
5. DB00399 (Zoledronic acid) is a drug used to prevent skeletal fractures in patients with cancers such as multiple myeloma and prostate cancer, and some studies show interesting links with leukemia (31).
6. DB00773 (Etoposide) is used for first-line treatment in patients with small-cell lung cancer. It is also used to treat other malignancies such as lymphoma or nonlymphocytic leukemia, according to DrugBank.
7. DB01254 (Dasatinib) is used in patients with Chronic Myelogenous leukemia, according to DrugBank.
8. DB01268 (Sunitinib): A phase I/II study of sunitinib and intensive chemotherapy for AML patients with FLT3 mutations has been achieved with encouraging results (32).
9. DB06589 (Pazopanib): This drug is in clinical trial phase II for patients with AML (33).
10. DB00530 (Erlotinib) is an inhibitor of the epidermal growth factor receptor (EGFR) tyrosine kinase that is used in the treatment of several types of cancer, including leukemia (34).

**Table S3.** MultiXrank parameters used for node prioritisation in leukemia.

| $r$ | $\delta$ | $\tau$ | $\lambda$ | $\eta$ |
|---|---|---|---|---|
| 0.7 | $\begin{bmatrix} 1/2 & 1/2 \end{bmatrix}$ | $\begin{bmatrix} 1/3 & 1/3 & 1/3 & 0 \\ 1/4 & 1/4 & 1/4 & 1/4 \end{bmatrix}$ | $\begin{bmatrix} 1/2 & 1/2 \\ 1/2 & 1/2 \end{bmatrix}$ | $\begin{bmatrix} 1/2 & 1/2 \end{bmatrix}$ |

**Table S4.** Top 10 prioritised genes for leukemia, associated MultiXrank scores, degree and distance to seed nodes.

| Genes | Scores | Degree | Distance to HRAS | Distance to Tipifarnib |
|---|---|---|---|---|
| CYP3A4 | 0.010934 | 5 | 3 | 1 |
| FNTB | 0.010690 | 16 | 3 | 1 |
| RAF1 | 0.001010 | 155 | 1 | 2 |
| RASGRP1 | 0.000984 | 142 | 1 | 3 |
| DGKZ | 0.000780 | 17 | 1 | 4 |
| RIN1 | 0.000746 | 55 | 1 | 3 |
| BRAF | 0.000237 | 168 | 1 | 2 |
| RASA1 | 0.000215 | 83 | 1 | 3 |
| AURKA | 0.000207 | 178 | 1 | 2 |
| ARAF | 0.000205 | 54 | 1 | 3 |

**Table S5.** Top 10 prioritised drugs for leukemia, associated MultiXrank scores, degree and distance to seed nodes.

| Drugs | Scores | Degree | Distance to HRAS | Distance to Tipifarnib |
|---|---|---|---|---|
| DB00637 | 0.000357 | 26 | 2 | 2 |
| DB01380 | 0.000349 | 69 | 4 | 2 |
| DB00630 | 0.000326 | 32 | 4 | 2 |
| DB00398 | 0.000267 | 108 | 2 | 1 |
| DB00399 | 0.000195 | 24 | 3 | 1 |
| DB00773 | 0.000191 | 80 | 3 | 1 |
| DB01254 | 0.000163 | 435 | 2 | 2 |
| DB01268 | 0.000144 | 157 | 1 | 2 |
| DB06589 | 0.000100 | 171 | 2 | 3 |
| DB00530 | 0.000085 | 95 | 2 | 2 |

#### B. Node prioritisation in Epilepsy.

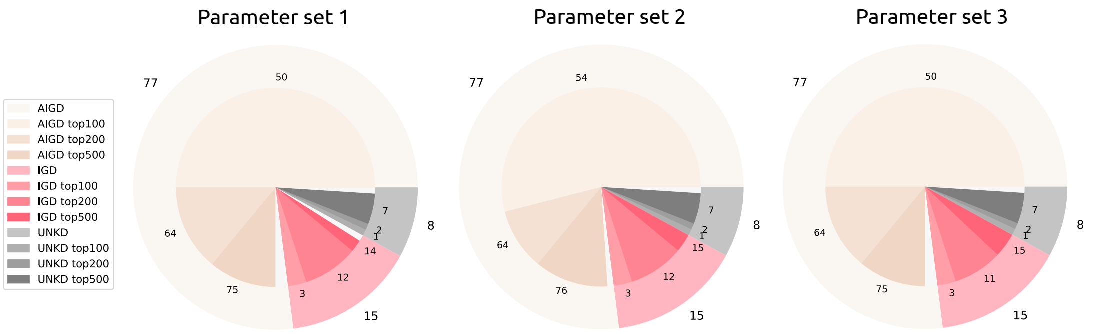

**Fig. S4.** Ictogenic properties of the 100 drugs prioritised by Hetionet and overlap with the top 100, 200, and 500 drugs prioritised by MultiXrank (parameter set 1 to 3, see Supplementary Table S6). The data on ictogenic properties has been sourced from the Hetionet study. AIGD are anti-ictogenic drugs that have a seizure suppressor effect (beige), IGD are ictogenic drugs (pink), and UKND are drugs with unknown effects (grey). The outer circle of the pie chart displays the number of AIGD, IGD, and UNKD drugs amongst the 100 drugs prioritised by Hetionet. The center of the pie chart displays the number of drugs from each category that were also prioritised by MultiXrank in the top-100 (displayed in lighter shades), top-200 (displayed in middle shade), and top-500 drugs (displayed in darker shades). The drugs prioritised in the top-500 include the drugs prioritised in the top-200, and the top-200 includes the drugs prioritised in the top-100. The white part of the pie chart corresponds to the drugs prioritised by Hetionet that are not in our MultiXrank top-500 prioritised drugs.

**Table S6.** Four sets of MultiXrank parameters used for node prioritisation in epilepsy. It is to note that only the $\lambda$ parameter differs.

| $r$ | $\delta$ | $\tau$ | $\lambda$ | $\eta$ |
|---|---|---|---|---|
| 0.7 | $\begin{bmatrix} 1/2 & 0 & 0 & 0 & 0 & 0 & 0 & 0 & 0 \end{bmatrix}$ | $\begin{bmatrix} 1/3 & 1/3 & 1/3 & 0 \\ 1 & 0 & 0 & 0 \\ 1 & 0 & 0 & 0 \\ 1 & 0 & 0 & 0 \\ 1 & 0 & 0 & 0 \\ 1 & 0 & 0 & 0 \\ 1 & 0 & 0 & 0 \\ 1 & 0 & 0 & 0 \\ 1 & 0 & 0 & 0 \end{bmatrix}$ | $\begin{bmatrix} 1/9 & 1/9 & 1/9 & 1/9 & 1/9 & 1/9 & 1/9 & 1/9 & 1/9 \\ 1/9 & 1/9 & 1/9 & 1/9 & 1/9 & 1/9 & 1/9 & 1/9 & 1/9 \\ 1/9 & 1/9 & 1/9 & 1/9 & 1/9 & 1/9 & 1/9 & 1/9 & 1/9 \\ 1/9 & 1/9 & 1/9 & 1/9 & 1/9 & 1/9 & 1/9 & 1/9 & 1/9 \\ 1/9 & 1/9 & 1/9 & 1/9 & 1/9 & 1/9 & 1/9 & 1/9 & 1/9 \\ 1/9 & 1/9 & 1/9 & 1/9 & 1/9 & 1/9 & 1/9 & 1/9 & 1/9 \\ 1/9 & 1/9 & 1/9 & 1/9 & 1/9 & 1/9 & 1/9 & 1/9 & 1/9 \\ 1/9 & 1/9 & 1/9 & 1/9 & 1/9 & 1/9 & 1/9 & 1/9 & 1/9 \\ 1/9 & 1/9 & 1/9 & 1/9 & 1/9 & 1/9 & 1/9 & 1/9 & 1/9 \end{bmatrix}$ | $\begin{bmatrix} 0 & 1 & 0 & 0 & 0 & 0 & 0 & 0 & 0 \end{bmatrix}$ |

Parameter set 1

| $r$ | $\delta$ | $\tau$ | $\lambda$ | $\eta$ |
|---|---|---|---|---|
| 0.7 | $\begin{bmatrix} 1/2 & 0 & 0 & 0 & 0 & 0 & 0 & 0 & 0 \end{bmatrix}$ | $\begin{bmatrix} 1/3 & 1/3 & 1/3 & 0 \\ 1 & 0 & 0 & 0 \\ 1 & 0 & 0 & 0 \\ 1 & 0 & 0 & 0 \\ 1 & 0 & 0 & 0 \\ 1 & 0 & 0 & 0 \\ 1 & 0 & 0 & 0 \\ 1 & 0 & 0 & 0 \\ 1 & 0 & 0 & 0 \end{bmatrix}$ | $\begin{bmatrix} 0 & 1/9 & 1/9 & 1/9 & 1/9 & 1/9 & 1/9 & 1/9 & 1/9 \\ 1/9 & 0 & 1/9 & 1/9 & 1/9 & 1/9 & 1/9 & 1/9 & 1/9 \\ 2/9 & 2/9 & 1/9 & 2/9 & 2/9 & 2/9 & 2/9 & 2/9 & 2/9 \\ 1/9 & 1/9 & 1/9 & 0 & 1/9 & 1/9 & 1/9 & 1/9 & 1/9 \\ 1/9 & 1/9 & 1/9 & 1/9 & 0 & 1/9 & 1/9 & 1/9 & 1/9 \\ 1/9 & 1/9 & 1/9 & 1/9 & 1/9 & 0 & 1/9 & 1/9 & 1/9 \\ 1/9 & 1/9 & 1/9 & 1/9 & 1/9 & 1/9 & 0 & 1/9 & 1/9 \\ 1/9 & 1/9 & 1/9 & 1/9 & 1/9 & 1/9 & 1/9 & 0 & 1/9 \\ 1/9 & 1/9 & 1/9 & 1/9 & 1/9 & 1/9 & 1/9 & 1/9 & 0 \end{bmatrix}$ | $\begin{bmatrix} 0 & 1 & 0 & 0 & 0 & 0 & 0 & 0 & 0 \end{bmatrix}$ |

Parameter set 2

| $r$ | $\delta$ | $\tau$ | $\lambda$ | $\eta$ |
|---|---|---|---|---|
| 0.7 | $\begin{bmatrix} 1/2 & 0 & 0 & 0 & 0 & 0 & 0 & 0 & 0 \end{bmatrix}$ | $\begin{bmatrix} 1/3 & 1/3 & 1/3 & 0 \\ 1 & 0 & 0 & 0 \\ 1 & 0 & 0 & 0 \\ 1 & 0 & 0 & 0 \\ 1 & 0 & 0 & 0 \\ 1 & 0 & 0 & 0 \\ 1 & 0 & 0 & 0 \\ 1 & 0 & 0 & 0 \\ 1 & 0 & 0 & 0 \end{bmatrix}$ | $\begin{bmatrix} 0 & 1/18 & 1/18 & 1/18 & 1/18 & 1/18 & 1/18 & 1/18 & 1/18 \\ 1/9 & 1/9 & 1/9 & 1/9 & 1/9 & 1/9 & 1/9 & 1/9 & 1/9 \\ 2/9 & 3/18 & 3/18 & 5/18 & 5/18 & 5/18 & 5/18 & 5/18 & 5/18 \\ 1/9 & 1/9 & 1/9 & 0 & 1/9 & 1/9 & 1/9 & 1/9 & 1/9 \\ 1/9 & 1/9 & 1/9 & 1/9 & 0 & 1/9 & 1/9 & 1/9 & 1/9 \\ 1/9 & 1/9 & 1/9 & 1/9 & 1/9 & 0 & 1/9 & 1/9 & 1/9 \\ 1/9 & 1/9 & 1/9 & 1/9 & 1/9 & 1/9 & 0 & 1/9 & 1/9 \\ 1/9 & 1/9 & 1/9 & 1/9 & 1/9 & 1/9 & 1/9 & 0 & 1/9 \\ 1/9 & 1/9 & 1/9 & 1/9 & 1/9 & 1/9 & 1/9 & 1/9 & 0 \end{bmatrix}$ | $\begin{bmatrix} 0 & 1 & 0 & 0 & 0 & 0 & 0 & 0 & 0 \end{bmatrix}$ |

Parameter set 3

| $r$ | $\delta$ | $\tau$ | $\lambda$ | $\eta$ |
|---|---|---|---|---|
| 0.7 | $\begin{bmatrix} 1/2 & 0 & 0 & 0 & 0 & 0 & 0 & 0 & 02 \end{bmatrix}$ | $\begin{bmatrix} 1/3 & 1/3 & 1/3 & 0 \\ 1 & 0 & 0 & 0 \\ 1 & 0 & 0 & 0 \\ 1 & 0 & 0 & 0 \\ 1 & 0 & 0 & 0 \\ 1 & 0 & 0 & 0 \\ 1 & 0 & 0 & 0 \\ 1 & 0 & 0 & 0 \\ 1 & 0 & 0 & 0 \end{bmatrix}$ | $\begin{bmatrix} 0 & 1/9 & 1/9 & 1/9 & 1/9 & 1/9 & 1/9 & 1/9 & 1/9 \\ 1/9 & 0 & 1/9 & 1/9 & 1/9 & 1/9 & 1/9 & 1/9 & 1/9 \\ 3/9 & 3/9 & 2/9 & 3/9 & 3/9 & 3/9 & 3/9 & 3/9 & 3/9 \\ 1/9 & 1/9 & 1/9 & 0 & 1/9 & 1/9 & 1/9 & 1/18 & 1/18 \\ 1/9 & 1/9 & 1/9 & 1/9 & 0 & 1/9 & 1/9 & 1/9 & 1/9 \\ 1/9 & 1/9 & 1/9 & 1/9 & 1/9 & 0 & 1/9 & 1/9 & 1/9 \\ 1/9 & 1/9 & 1/9 & 1/9 & 1/9 & 1/9 & 0 & 1/9 & 1/9 \\ 1/18 & 1/18 & 1/18 & 1/18 & 1/18 & 1/18 & 1/18 & 0 & 1/18 \\ 1/18 & 1/18 & 1/18 & 1/18 & 1/18 & 1/18 & 1/18 & 1/18 & 0 \end{bmatrix}$ | $\begin{bmatrix} 0 & 1 & 0 & 0 & 0 & 0 & 0 & 0 & 0 \end{bmatrix}$ |

Parameter set 4

**Table S7.** DrugBank Categories associated to at least 7 drugs from the 41 drugs prioritised by MultiXrank (parameter set number 4, top-100) that are not prioritised by Hetionet. The first column of the table corresponds to the DrugBank Category and the second column ($N$) corresponds to the number of drugs mapped to each category. It is to note that some drugs belong to more than one class.

| **Drugs class** | **N** | **Drugs class** | **N** | **Drugs class** | **N** |
|---|---|---|---|---|---|
| Cytochrome P-450 Substrates | 24 | Peripheral Nervous System Agents | 10 | Agents causing hyperkalemia | 8 |
| Agents that produce hypertension | 18 | Antimigraine Preparations | 10 | Cytochrome P-450 CYP2C19 Substrates | 7 |
| Central Nervous System Depressants | 18 | Cardiovascular Agents | 10 | Cytochrome P-450 CYP3A4 Inhibitors | 7 |
| Neurotransmitter Agents | 17 | Sensory System Agents | 10 | Enzyme Inhibitors | 7 |
| Analgesics | 17 | P-glycoprotein substrates | 9 | Biogenic Amines | 7 |
| Heterocyclic Compounds Fused-Ring | 15 | Drugs that are Mainly Renally Excreted | 9 | Biogenic Monoamines | 7 |
| Cytochrome P-450 Enzyme Inhibitors | 14 | Serotonin Modulators | 9 | Selective Serotonin 5-HT1 Receptor Agonists | 7 |
| Cytochrome P-450 CYP3A4 Substrates | 14 | Serotonin Receptor Agonists | 9 | Selective Serotonin Agonists | 7 |
| Amines | 12 | Serotonin 5-HT1 Receptor Agonists | 9 | Serotonin 1b Receptor Agonists | 7 |
| Cytochrome P-450 CYP1A2 Substrates | 12 | Cytochrome P-450 CYP2D6 Substrates | 9 | Serotonin 1d Receptor Agonists | 7 |
| Serotonergic Drugs | 11 | Cytochrome P-450 CYP2C9 Substrates | 8 | Triptans | 7 |
| Antidepressive Agents | 10 | Indoles | 8 | P-glycoprotein inhibitors | 7 |
| Serotonin Agents | 10 | Analgesics Non-Narcotic | 8 | | |

#### 3. Supervised prediction of gene-disease associations

The classifiers were trained on MultiXrank scores generated from exploring the multilayer network, taking as seeds the gene and disease nodes considered in the association. We detail the procedure bellow.

1. **Creation of the training dataset from DisGeNET v2.0 (2014) associations**
   Gene-disease associations were obtained from an outdated version of DisGeNET (v2.0, 2014). We filtered out associations with scores below 0,5 in order to remove associations that had insufficient support in the literature and obtained 1914 gene-disease associations. These associations constitute our positive dataset. On the other hand, we generated a negative dataset by randomly picking 3828 pairs of gene and disease nodes that are not considered associated in DisGeNET v2.0 (2014). We selected twice as many negative associations than positive associations in the training set in order to account for class imbalance, which is likely to occur in real settings. We thus obtained 5742 labelled associations. We further split this dataset into a training and test set, keeping 30% of the associations for evaluating the performance of the models to predict DisGeNET v2.0 (2014) associations.

2. **Running MultiXrank for all associations**
   For each positive and negative gene-disease association defined in the training dataset, we used both the gene and the disease nodes as seeds when running MultiXrank. For positive associations, we also removed the bipartite edge between the two seeds before running MultiXrank. For each pair of seed (i.e. positive or negative association), we ran MultiXrank on the multilayer network with the parameters defined in Supplementary Table S8. These scores encode the similarity of all the nodes of the multilayer network with respect to the seeds, which we can use to train a classifier for predicting gene-disease associations.

3. **Training binary classifiers from MultiXrank scores**
   The output scores obtained with MultiXrank using the positive and negative gene-disease associations as seeds were used to train Random Forest and XGBoost classifiers. For both classifiers, we used various class weights, which are often used in imbalanced classifications problems for penalising classification errors on the minority class. The class weights used to train the classifiers are described in Supplementary Table S9, in which we also report the performances of each models in predicting the associations from DisGeNET v2.0 (2014) that were kept as a test set.

4. **Creation of the test dataset from DisGeNET v7.0 (2020) associations**
   To evaluate weather novel associations from DisGeNET v7.0 (2020) could have been predicted based on MultiXrank scores obtained using DisGeNET v2.0 (2014) associations only, we create a test dataset containing positive and negative gene-disease associations from DisGeNET v7.0 (2020). The positive associations were extracted from DisGeNET v7.0 (selecting associations above a threshold equal to 0.5) and the negative associations were generated by randomly selecting pairs of gene and disease that are not considered associated in DisGeNET v2.0. We obtained 7218 novel positive gene-disease associations, and randomly selected twice as many (i.e. 14 436) negative associations. We made sure all selected positive and negative associations were considering genes and diseases that were already present in the gene and disease layers of the multilayer network.

5. **Evaluation of the random forest binary classifier on the test dataset from DisGeNET v7.0 (2020) associations**
   We ran MultiXrank using as seeds the gene and disease nodes of each positive and negative gene-disease association of the test dataset, using the same parameters used for obtaining MultiXrank scores for the training dataset (Supplementary Table S8). We used MultiXrank output scores as input of the previously computed Random Forest and XGBoost models (trained on the data obtained with the gene-disease association of DisGeNET v2.0 (2014)) and predicted their label. Finally, we compared the predicted labels to the true labels. We report the results for each model in Supplementary Table S10.

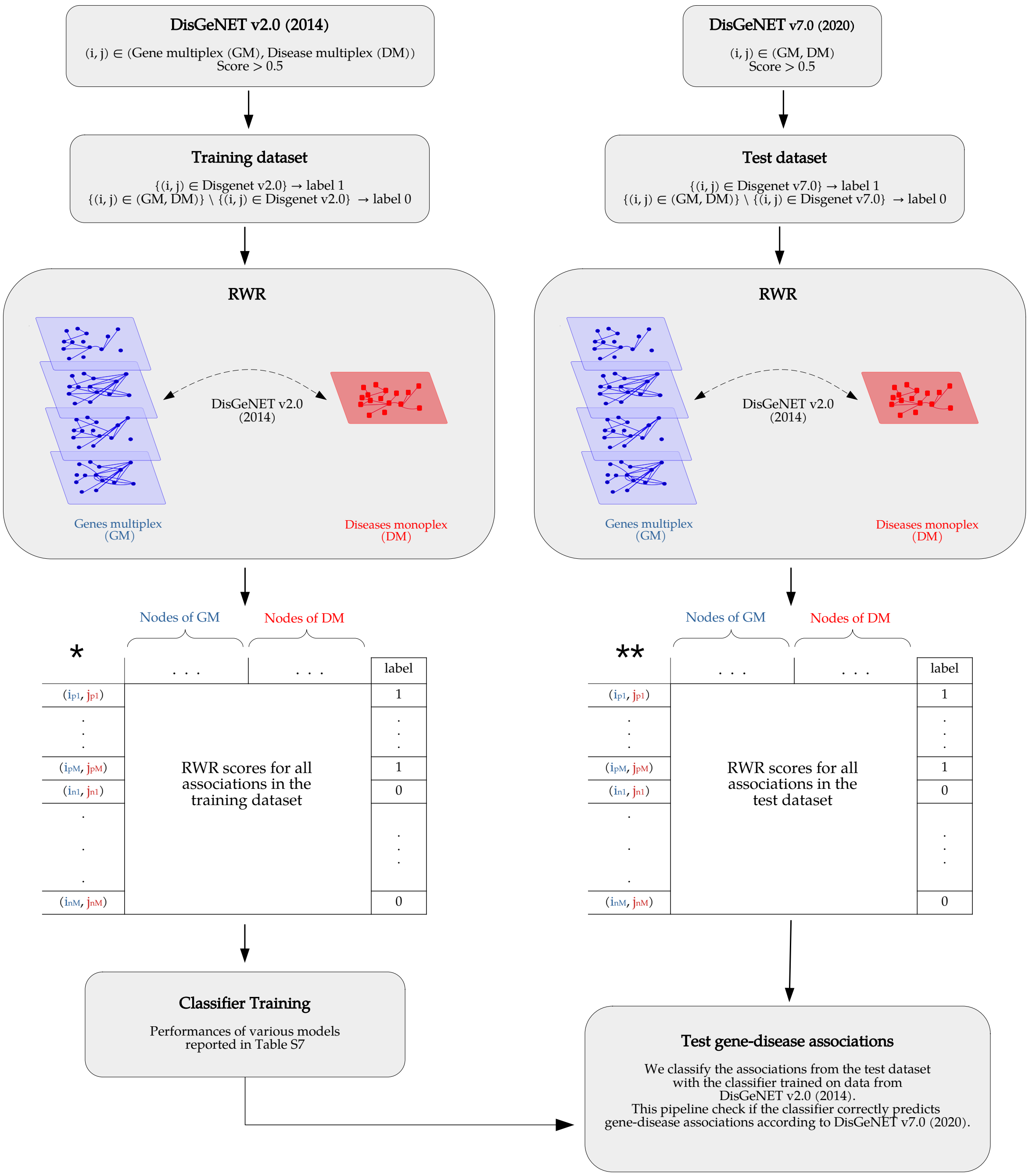

**Fig. S5.** Workflow of the random forest binary classifier. The left panel represents the training step. Here, the bipartite network connecting the gene multiplex and the disease monoplex network is built from DisGeNET v2.0 (2014). The gene-disease associations in DisGeNET v2.0 (2014) are also considered positive gene-disease associations. Negative gene-disease associations are sampled randomly. Then, MultiXrank is run using the gene and the disease nodes of each positive and negative gene-disease association, and the output scores are saved as described in the matrix *. This matrix is used with the positive and negative labels to train a random forest binary classifier. The right panel represents the test step. In this case, the true positive gene-disease associations are created from DisGeNET v7.0 (2020) and true negative gene-disease associations are sampled randomly. We next ran MultiXrank using as seeds the gene and disease nodes from each positive and negative gene-disease association, and saved the output scores in the matrix **. Finally, this ** matrix is used as an input of the previously trained random forest classifier to predict the labels of each gene-disease association. The predicted labels are then compared with the true known labels. The protocol is the same for two or three multiplex networks.

**Table S8.** MultiXrank parameters used for the supervised prediction of gene-disease associations.

| $r$ | $\delta$ | $\tau$ | $\lambda$ | $\eta$ |
|---|---|---|---|---|
| $0.7$ | $\begin{bmatrix} 1/2 & 0 \end{bmatrix}$ | $\begin{bmatrix} 1/3 & 1/3 & 1/3 \\ 1 & 0 & 0 \end{bmatrix}$ | $\begin{bmatrix} 1/2 & 1/2 \\ 1/2 & 1/2 \end{bmatrix}$ | $\begin{bmatrix} 1/2 & 1/2 \end{bmatrix}$ |

**Table S9.** XGBoost and Random Forest models and performance metrics for predicting Gene-Disease associations from DisGeNET v2.0 (2014). The models are trained and tested according to the protocol described in Supplementary Figure S5. We trained the XGBoost and Random Forest models using various class weights, large weights penalising classification errors for the corresponding class (pos., for positive associations and neg., for negative associations). We report several performance metrics for each model: Balanced Accuracy (defined as the average of recall obtained on each class), F1-score and number of True Positives (TP), False Positives (FP), True Negatives (TN) and False Negative (FN). The best performing model is highlighted in grey.

| Classifier | Class Weight Neg. | Class Weight Pos. | Balanced Accuracy | F1 Score | TP | FP | TN | FN |
|---|---|---|---|---|---|---|---|---|
| XGBoost | 0.1 | 0.9 | 0.853 | 0.792 | 508 | 197 | 948 | 70 |
| XGBoost | 0.2 | 0.8 | 0.844 | 0.787 | 481 | 164 | 981 | 97 |
| XGBoost | 0.3 | 0.7 | 0.848 | 0.795 | 472 | 138 | 1007 | 106 |
| XGBoost | 0.4 | 0.6 | 0.838 | 0.785 | 452 | 121 | 1024 | 126 |
| XGBoost | 0.5 | 0.5 | 0.822 | 0.768 | 421 | 97 | 1048 | 157 |
| XGBoost | 0.6 | 0.4 | 0.816 | 0.762 | 412 | 92 | 1053 | 166 |
| XGBoost | 0.7 | 0.3 | 0.807 | 0.751 | 390 | 70 | 1075 | 188 |
| XGBoost | 0.8 | 0.2 | 0.801 | 0.745 | 377 | 57 | 1088 | 201 |
| XGBoost | 0.9 | 0.1 | 0.767 | 0.695 | 331 | 44 | 1101 | 247 |
| Random Forest | 0.1 | 0.9 | 0.740 | 0.651 | 320 | 85 | 1060 | 258 |
| Random Forest | 0.2 | 0.8 | 0.749 | 0.666 | 319 | 61 | 1084 | 259 |
| Random Forest | 0.3 | 0.7 | 0.739 | 0.649 | 312 | 71 | 1074 | 266 |
| Random Forest | 0.4 | 0.6 | 0.772 | 0.701 | 347 | 65 | 1080 | 231 |
| Random Forest | 0.5 | 0.5 | 0.769 | 0.697 | 348 | 73 | 1072 | 230 |
| Random Forest | 0.6 | 0.4 | 0.784 | 0.718 | 369 | 81 | 1064 | 209 |
| Random Forest | 0.7 | 0.3 | 0.792 | 0.729 | 386 | 95 | 1050 | 192 |
| Random Forest | 0.8 | 0.2 | 0.812 | 0.756 | 411 | 99 | 1046 | 167 |
| Random Forest | 0.9 | 0.1 | 0.818 | 0.762 | 422 | 108 | 1037 | 156 |

**Table S10.** XGBoost and Random Forest models and performance metrics for predicting Gene-Disease associations from DisGeNET v7.0 (2020). The models are the same as those presented in Supplementary Table S9, and have not been retrained for predicting DisGeNET v7.0 (2020) associations. We report several performance metrics for each model: Balanced Accuracy (defined as the average of recall obtained on each class), F1-score and number of True Positives (TP), False Positives (FP), True Negatives (TN) and False Negative (FN). The best performing model is highlighted in grey.

| Model | Class Weight Neg. | Class Weight Pos. | Balanced Accuracy | F1 Score | TP | FP | TN | FN |
|---|---|---|---|---|---|---|---|---|
| XGBoost | 0.1 | 0.9 | 0.642 | 0.530 | 4026 | 3950 | 10486 | 3192 |
| XGBoost | 0.2 | 0.8 | 0.633 | 0.509 | 3631 | 3423 | 11013 | 3587 |
| XGBoost | 0.3 | 0.7 | 0.633 | 0.501 | 3402 | 2972 | 11464 | 3816 |
| XGBoost | 0.4 | 0.6 | 0.627 | 0.484 | 3140 | 2613 | 11823 | 4078 |
| XGBoost | 0.5 | 0.5 | 0.622 | 0.468 | 2901 | 2274 | 12162 | 4317 |
| XGBoost | 0.6 | 0.4 | 0.616 | 0.449 | 2658 | 1971 | 12465 | 4560 |
| XGBoost | 0.7 | 0.3 | 0.613 | 0.438 | 2525 | 1785 | 12651 | 4693 |
| XGBoost | 0.8 | 0.2 | 0.606 | 0.415 | 2279 | 1498 | 12938 | 4939 |
| XGBoost | 0.9 | 0.1 | 0.598 | 0.388 | 2030 | 1218 | 13218 | 5188 |
| Random Forest | 0.1 | 0.9 | 0.566 | 0.352 | 1975 | 2035 | 12401 | 5243 |
| Random Forest | 0.2 | 0.8 | 0.576 | 0.361 | 1972 | 1745 | 12691 | 5246 |
| Random Forest | 0.3 | 0.7 | 0.578 | 0.367 | 2029 | 1817 | 12619 | 5189 |
| Random Forest | 0.4 | 0.6 | 0.583 | 0.381 | 2155 | 1925 | 12511 | 5063 |
| Random Forest | 0.5 | 0.5 | 0.594 | 0.404 | 2309 | 1914 | 12522 | 4909 |
| Random Forest | 0.6 | 0.4 | 0.600 | 0.422 | 2493 | 2102 | 12334 | 4725 |
| Random Forest | 0.7 | 0.3 | 0.609 | 0.446 | 2732 | 2307 | 12129 | 4486 |
| Random Forest | 0.8 | 0.2 | 0.615 | 0.462 | 2932 | 2556 | 11880 | 4286 |
| Random Forest | 0.9 | 0.1 | 0.618 | 0.474 | 3129 | 2847 | 11589 | 4089 |

### 4. Diffusion profiles comparison to unveil immune diseases similarities

#### A. PCHi-C and TAD experiments reflects the tree lineage of hematopoietic cells.

Before subjecting each hematopoietic cell line-specific multilayer network to analysis using MultiXrank our objective was to ascertain whether the PCHi-C and TAD datasets could effectively distinguish between the different hematopoietic cell lines. The hierarchical lineage of hematopoietic cells is visually represented in Supplementary Figure S1. The tree lineage is composed of two branches: the first branch encompasses lymphoid cells, while the second branch encompasses myeloid cells.

The lymphoid cells branch is further divided into two sub-branches: the first one contains the B-cells with the naive B cells (nB), and the second one contains the T-cells with the Naive CD4+ T cells (nCD4) and the Naive CD8+ T cells (nCD8).

The myeloid cells branch is also further divided into two other branches: the first one contains the Monocytes (Mon) and the Neutrophils (Neu), and the second one contains the Megakaryocytes (MK), and the Erythroblasts (Ery). The Macrophages M0 (Mac0) are differentiated from monocytes (35).

To assess the ability of the PCHi-C and TAD datasets to differentiate among these diverse cell types, we first defined a similarity measure to evaluate the similarity of cell lines according to the PCHi-C and TAD datasets. We used the Jaccard index as a similarity metric, to evaluate the proportion of shared PCHi-C fragments and TAD between hematopoietic cells:

$$J_{i,j} = \frac{D_i \cap D_j}{D_i \cup D_j} \quad [1]$$

where $D_i$ and $D_j$ the datasets (PCHi-C or TAD) corresponding to the hematopoietic cells $i$ and $j$.

We projected the obtained similarity matrix into a 2D Principal Component Analysis (PCA) space (Supplementary Figure S6).

We can discern a distinct division between lymphoid cells (represented by red dots) and myeloid cells (represented by blue dots) in the PCHi-C fragment PCA and, to a lesser extent, in the TAD PCA. Furthermore,

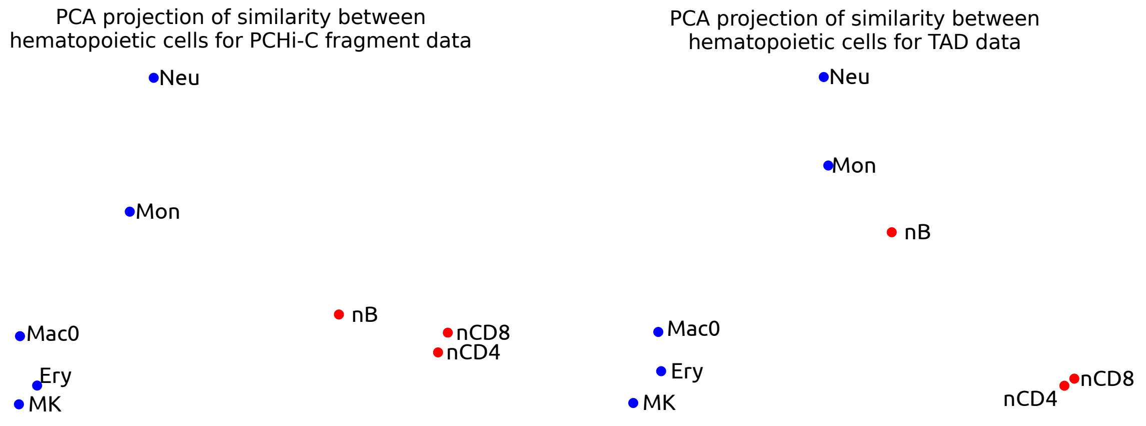

**Fig. S6.** 2D PCA projection of the Jaccard index similarities between the different hematopoietic cell types. Left: similarities computed on the PCHi-C fragment dataset. Right: similarities computed on the TAD dataset. Red: Lymphoid cells. Blue: Myeloid cells.

on both PCA plots, there is proximity between nCD4 and nCD8 cells, while nB cells are positioned closely to both of these hematopoietic cell types. The Mon and the Neu cell types are also closer to each other than to the other cell types. Finally, the Ery and MK show good proximity in both the PCHi-C fragment and the TAD PCA.

In conclusion, those projections show that both PCHi-C and TAD datasets reflect the tree lineage of the hematopoietic cells.

#### B. RWR exploration of the multilayer networks including PCHi-C and TAD network layers reflects the tree lineage of the hematopoietic cells.

In the previous section, we demonstrated that the PCHi-C fragments and TAD datasets can effectively capture the tree lineage of hematopoietic cells. We now aim to illustrate that MultiXrank immune diseases diffusion profiles computed on the height hematopoietic multilayer networks, can also capture this lineage.

We used MultiXrank to explore the hematopoietic multilayer networks, using as seeds the 131 different immune diseases iteratively (Supplementary Table S13). Hence, we produced 131 different RWR output scores (one for each immune disease) for each of the 8 hematopoietic multilayer network. In addition, since MultiXrank produces output scores for each of the 4 node types in the multilayer networks (gene/protein nodes, disease nodes, PCHI-C fragment nodes and TAD nodes), we obtained overall 131 (immune disease seeds) $\times$ 8 (cell types) $\times$ 4 (node types) RWR vectors.

Based on these RWR vectors, we defined a similarity measure between the different hematopoietic cell types. First, we compute a disease-disease distance matrix for each node type and hematopoietic network independently. This is done by comparing the rankings of nodes obtained for every pair of immune disease seed (equation 1, Materials and Methods).

To compute cell-cell similarities from those disease-disease distance matrices, we compute the Pearson's correlations coefficients between each disease vector for a specific node type and average it, as detailed below:

$$P^t_{cell_i,cell_j} = \frac{\sum_{d=1}^{N_d} cor(D_d^{cell_i,t}, D_d^{cell_j,t})}{N_d}$$

with $P^t_{cell_i,cell_j}$ the averaged Pearson's correlations for hematopoietic cells $i$ and $j$ for node type $t$, $N_d$ the total number of diseases, $D_d^{cell_i,t}$ the distance vector of disease $d$ in cell $cell_i$ and node type $t$, and $D_d^{cell_j,t}$ the distance vector of disease $d$ in cell $cell_j$ and node type $t$.

This produces one similarity matrix of size $8 \times 8$ for each node type (i.e. protein, disease, PCHi-C fragment and TAD) independently.

We projected those 4 node types similarity matrices into a 2D PCA space (Supplementary Figure S7). Those result show that the scores obtained on TAD nodes were the ones that best captured the tree lineage of hematopoietic cells, although the disease and PCHi-C fragment node types have also captured some similarities between nearby cells in the tree lineage. Scores obtained on the protein node type did not allow us to recover the lineage.

Finally, we fuse the cell-cell similarity matrices from the four node types by computing their average Pearson correlations across node types as follow:

$$P^{int}_{cell_i, cell_j} = \frac{\sum_{t=1}^{N_t} cor(P^t_{cell_i}, P^t_{cell_j})}{N_t}$$

with $P^{int}_{i,j}$ the integrated averaged Pearson's correlations for hematopoietic cell $cell_i$ and $cell_j$, $N_t$ the total number of node types in the multilayer network, and with $P^t_i$ (resp. $P^t_j$) the correlation vector of cell $cell_i$ (resp. $cell_j$) computed for node type $t$.

The matrix obtained from this procedure represents the similarity of hematopoietic cells based on the integrated MultiXrank scores. We again projected the matrix into a 2D PCA space (Supplementary Figure S8). We observe that the integrated MultiXrank score accurately mirrors the lineage of hematopoietic cells. Indeed, lymphoid cells (red) and myeloid cells (blue) exhibit a separation in the PCA space (PC1). Moreover, nCD4 and nCD8 cells are in close proximity to each other and relative proximity with nB cells. Furthermore, Macrophage (Mac0) and its precursor, Monocyte (Mon), appear closely situated. Additionally, Erythrocyte (Ery) and Megakaryocyte (MK), both stemming from the same progenitor, also exhibit high proximity to each other.

This projection is the one that most effectively summarises the tree lineage, when compared to projecting the similarity matrices of the four node types independently. It demonstrates that MultiXrank is able to capture meaning-full patterns in a multilayer network and reinforces the importance of incorporating different layers in the hematopoietic multilayer networks.

**Table S11.** MultiXrank parameters used for diffusion profiles comparison to unveil immune diseases similarities.

| $r$ | $\delta$ | $\tau$ | $\lambda$ | $\eta$ |
|---|---|---|---|---|
| 0.7 | $\begin{bmatrix} 1/2 & 0 & 0 & 0 \end{bmatrix}$ | $\begin{bmatrix} 1/3 & 1/3 & 1/3 \\ 1 & 0 & 0 \\ 1 & 0 & 0 \\ 1 & 0 & 0 \end{bmatrix}$ | $\begin{bmatrix} 1/4 & 1/4 & 1/4 & 1/4 \\ 1/4 & 1/4 & 1/4 & 1/4 \\ 1/4 & 1/4 & 1/4 & 1/4 \\ 1/4 & 1/4 & 1/4 & 1/4 \end{bmatrix}$ | $\begin{bmatrix} 0 & 0 & 0 & 1 \end{bmatrix}$ |

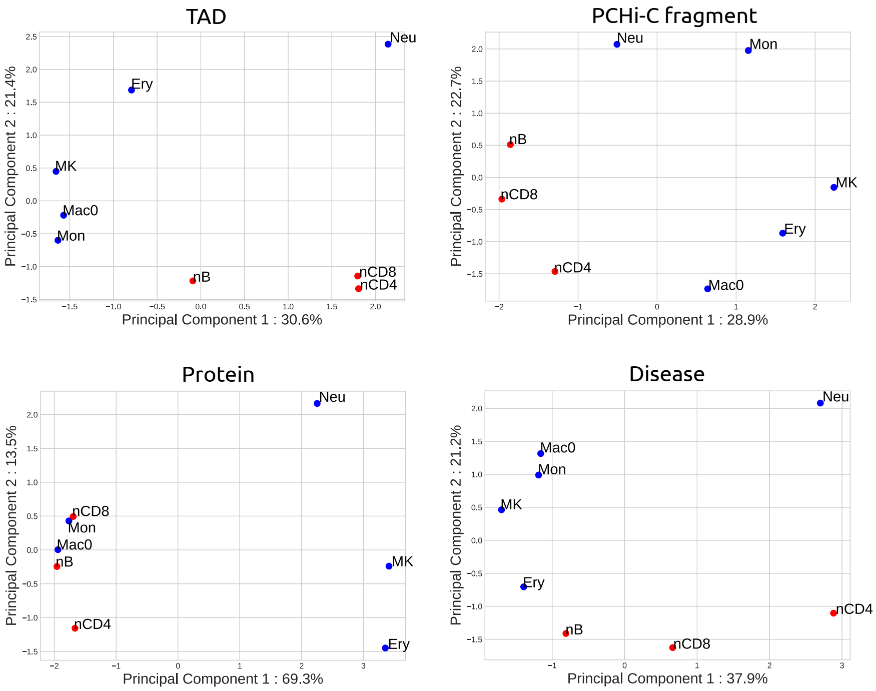

**Fig. S7.** 2D PCA projection of the similarity between the hematopoietic cell types. The tree lineage of hematopoietic cells is correctly found with the PCHi-C fragment, TAD and disease output scores of MultiXrank. However, the tree lineage is not recovered for the protein output scores.

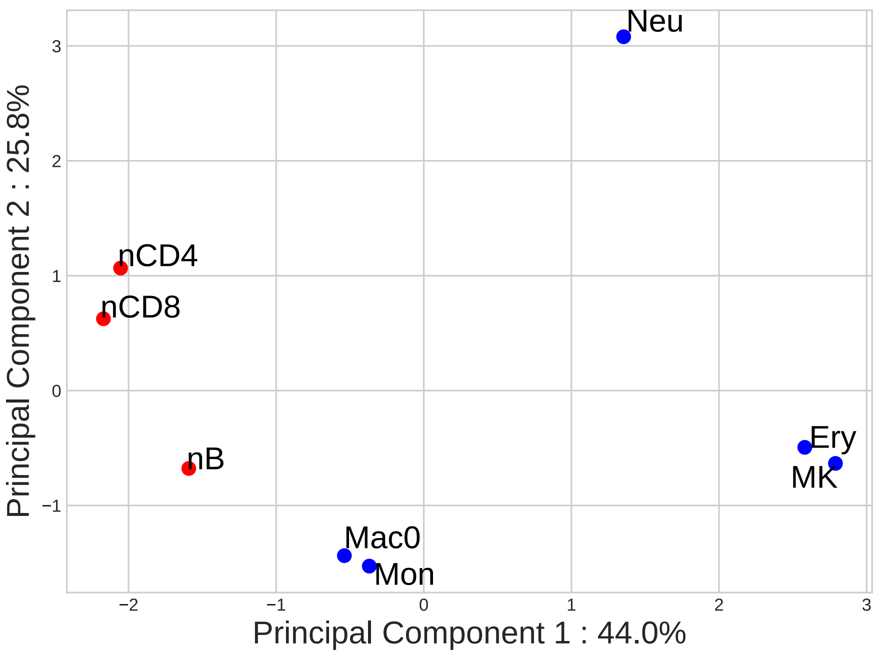

**Fig. S8.** 2D PCA projection of hematopoietic cell similarities with respect to the integrated MultiXrank output scores effectively visualises the similarity between various hematopoietic cell types. In this projection, lymphoid cells (depicted in red) and myeloid cells (depicted in blue) exhibit a relative separation within the PCA space. Moreover, nCD4 and nCD8 cells are in close proximity to each other and relative proximity with nB cells. Furthermore, Macrophage (Mac0) and its precursor, Monocyte (Mon), appear closely situated. Additionally, Erythrocyte (Ery) and Megakaryocyte (MK), both stemming from the same progenitor, also exhibit high proximity to each other.

**Table S12.** Composition and characteristics of the immune disease clusters

| Cl | Diseases | Characteristics |
|---|---|---|
| **0** | Amyloidosis familial visceral ; Autoimmune polyglandular syndrome type 1 ; Autosomal recessive early-onset inflammatory bowel disease 25 ; Autosomal recessive early-onset inflammatory bowel disease 28 ; Behcet's disease ; Blau syndrome ; Chronic recurrent multifocal osteomyelitis (CRMO) ; Cicatricial pemphigoid ; Cryoglobulinemic vasculitis ; Eosinophilic esophagitis (EoE) ; Eosinophilic fasciitis ; Familial cold autoinflammatory syndrome ; Familial Mediterranean fever ; Giant cell arteritis (temporal arteritis) ; Goodpasture's syndrome ; Granulomatosis with Polyangiitis ; Hyper-IgD syndrome ; Kawasaki disease (Mucocutaneous Lymph Node Syndrome) ; Majeed syndrome ; Myasthenia gravis ; Mycosis fungoides ; Myositis ; Netherton syndrome ; Papillon Lefevre syndrome ; Primary biliary cirrhosis ; Pyogenic arthritis, pyoderma gangrenosum and acne ; Rheumatoid arthritis ; Schmidt syndrome ; Takayasu's arteritis ; Tumor necrosis factor receptor-associated periodic syndrome ; Type 1 diabetes | Inflammatory, Autoinflammatory and Autoimmune Diseases. |
| **1** | Activated PI3K-Delta Syndrome 1 ; Activated PI3K-Delta Syndrome 2 ; Agammaglobulinemia ; Autoimmune lymphoproliferative syndrome ; Autoimmune lymphoproliferative syndrome due to CTLA4 haploinsuffiency ; Autosomal dominant hyper IgE syndrome ; Autosomal recessive candidiasis familial chronic mucocutaneous ; C1q deficiency ; Combined immunodeficiency with skin granulomas ; Complement component 2 deficiency ; Complement component 8 deficiency type 1 ; Complement component 8 deficiency type 2 ; Cyclic neutropenia ; Deficiency of interleukin-1 receptor antagonist ; Dendritic cell, monocyte, B lymphocyte, and natural killer lymphocyte deficiency ; Familial hemophagocytic lymphohistiocytosis ; Felty's syndrome ; Hashimoto's thyroiditis ; Hepatic venoocclusive disease with immunodeficiency ; Hypogammaglobulinemia AGM2 (IGLL1) ; Hypogammaglobulinemia AGM3 (CD79A) ; Hypogammaglobulinemia AGM4 (BLNK) ; Hypogammaglobulinemia AGM5 (LRRC8A) ; Hypogammaglobulinemia AGM6 (CD79B) ; ICF syndrome ; IL12RB1 deficiency ; Immune dysfunction with T-cell inactivation due to calcium entry defect 1 ; Immune dysregulation, polyendocrinopathy and enteropathy X-linked ; Immune thrombocytopenic purpura (ITP) ; Immunodeficiency with hyper IgM type 1 ; Immunodeficiency with hyper IgM type 2 ; Immunodeficiency with hyper IgM type 3 ; Immunodeficiency with hyper IgM type 4 ; Immunodeficiency with hyper IgM type 5 ; Immunodeficiency without anhidrotic ectodermal dysplasia ; Immunoglobulin A deficiency 2 ; Intestinal atresia multiple ; IRAK-4 deficiency ; Isolated growth hormone deficiency type 3 ; Leukocyte adhesion deficiency type 1 ; LRBA deficiency ; MHC class 1 deficiency ; MYD88 deficiency ; Neutrophil-specific granule deficiency ; PGM3-CDG ; Poikiloderma with neutropenia ; Purine nucleoside phosphorylase deficiency ; Reticular dysgenesis ; Severe congenital neutropenia X-linked ; Short-limb skeletal dysplasia with severe combined immunodeficiency ; Sjögren's syndrome ; Spondyloenchondrodysplasia with immune dysregulation ; T-cell immunodeficiency, congenital alopecia and nail dystrophy ; Vici syndrome ; WHIM syndrome ; Wiskott Aldrich syndrome ; X-linked immunodeficiency with magnesium defect, Epstein-Barr virus infection and neoplasia ; X-linked lymphoproliferative syndrome ; ZAP-70 deficiency | Immunodeficiencies, Primary Immunodeficiencies, Increased Susceptibility to Infections |
| **2** | 22q11.2 deletion syndrome (DiGeorge syndrome) ; Acute myeloid leukemia ; Aicardi-Goutieres syndrome 1 ; Aicardi-Goutieres syndrome 2 ; Asymmetric crying face association ; Ataxia telangiectasia ; Barth syndrome ; Bloom syndrome ; Burkitt lymphoma ; Cartilage-hair hypoplasia ; CHARGE syndrome ; Chediak-Higashi syndrome ; Cherubism ; Chronic lymphocytic leukemia ; Chronic myelogenous leukemia ; Congenital heart block ; Epidermodysplasia verruciformis ; Glycogen storage disease type 1B ; Griscelli syndrome type 2 ; Hereditary folate malabsorption ; Hermansky Pudlak syndrome 2 ; Hodgkin's lymphoma ; Inclusion body myositis (IBM) ; Lichen sclerosus ; Melkersson-Rosenthal syndrome ; Meniere's disease ; Myelodysplastic syndromes ; Non-Hodgkin lymphoma ; Osteopetrosis autosomal recessive 7 ; Parry Romberg syndrome ; Pearson syndrome ; Pernicious anemia (PA) ; Pruritic urticarial papules plaques of pregnancy ; Pseudo-TORCH syndrome ; Raynaud's phenomenon ; Schimke immuno-osseous dysplasia ; Shprintzen syndrome ; Shwachman-Diamond syndrome ; Stiff person syndrome (SPS) ; TARP syndrome ; Woods Black Norbury syndrome | Blood-associated diseases (including Leukemia and Lymphoma), Cardiovascular, Dermatological, Hepatic, Muscular, Neurological, Neuromuscular and Skeletal diseases. |

**Table S13.** List of the $131$ immune diseases considered in this study. The first column represents the number used to identify each disease in the t-SNE projection. The second column represents the name of the disease. The third column is the UMLS identifier of the disease.

| ID | Immune Disease Name | UMLS |
|---|---|---|
| 0 | Agammaglobulinemia | UMLS:C0221026 |
| 1 | Behcet's disease | UMLS:C0004943 |
| 2 | Chronic recurrent multifocal osteomyelitis (CRMO) | UMLS:C0410422 |
| 3 | Cicatricial pemphigoid | UMLS:C1282359 |
| 4 | Congenital heart block | UMLS:C0149530 |
| 5 | Eosinophilic esophagitis (EoE) | UMLS:C0341106 |
| 6 | Eosinophilic fasciitis | UMLS:C0264005 |
| 7 | Giant cell arteritis (temporal arteritis) | UMLS:C1956391 |
| 8 | Goodpasture's syndrome | UMLS:C0403529 |
| 9 | Granulomatosis with Polyangiitis | UMLS:C3495801 |
| 10 | Hashimoto's thyroiditis | UMLS:C0677607 |
| 11 | Hypogammaglobulinemia AGM2 (IGLL1) | UMLS:C3150750 |
| 12 | Hypogammaglobulinemia AGM3 (CD79A) | UMLS:C3150751 |
| 13 | Hypogammaglobulinemia AGM4 (BLNK) | UMLS:C3150752 |
| 14 | Hypogammaglobulinemia AGM5 (LRRC8A) | UMLS:C3150753 |
| 15 | Hypogammaglobulinemia AGM6 (CD79B) | UMLS:C3150207 |
| 16 | Immune thrombocytopenic purpura (ITP) | UMLS:C0398650 |
| 17 | Inclusion body myositis (IBM) | UMLS:C0238190 |
| 18 | Kawasaki disease (Mucocutaneous Lymph Node Syndrome) | UMLS:C0026691 |
| 19 | Lichen sclerosus | UMLS:C0023652 |
| 20 | Meniere's disease | UMLS:C0025281 |
| 21 | Myasthenia gravis | UMLS:C0026896 |
| 22 | Myositis | UMLS:C0027121 |
| 23 | Parry Romberg syndrome | UMLS:C0015458 |
| 24 | Pernicious anemia (PA) | UMLS:C0002892 |
| 25 | Primary biliary cirrhosis | UMLS:C0008312 |
| 26 | Raynaud's phenomenon | UMLS:C0034734 |
| 27 | Rheumatoid arthritis | UMLS:C0003873 |
| 28 | Schmidt syndrome | UMLS:C0085860 |
| 29 | Sjögren's syndrome | UMLS:C1527336 |
| 30 | Stiff person syndrome (SPS) | UMLS:C0085292 |
| 31 | Takayasu's arteritis | UMLS:C0039263 |
| 32 | Burkitt lymphoma | UMLS:C0006413 |
| 33 | Acute myeloid leukemia | UMLS:C0023467 |
| 34 | Chronic lymphocytic leukemia | UMLS:C0023434 |
| 35 | Chronic myelogenous leukemia | UMLS:C0023473 |
| 36 | Hodgkin's lymphoma | UMLS:C0019829 |
| 37 | Myelodysplastic syndromes | UMLS:C3463824 |
| 38 | Non-Hodgkin lymphoma | UMLS:C0024305 |
| 39 | Mycosis fungoides | UMLS:C0026948 |
| 40 | Type 1 diabetes | UMLS:C0011854 |

| ID | Name | UMLS |
|---|---|---|
| 41 | Shprintzen syndrome | UMLS:C0220704 |
| 42 | Asymmetric crying face association | UMLS:C0431406 |
| 43 | 22q11.2 deletion syndrome (DiGeorge syndrome) | UMLS:C0012236 |
| 44 | Pseudo-TORCH syndrome | UMLS:C3489725 |
| 45 | Aicardi-Goutieres syndrome 1 | UMLS:C0796126 |
| 46 | Aicardi-Goutieres syndrome 2 | UMLS:C3489724 |
| 47 | Amyloidosis familial visceral | UMLS:C0268389 |
| 48 | Ataxia telangiectasia | UMLS:C0004135 |
| 49 | Autoimmune lymphoproliferative syndrome | UMLS:C1328840 |
| 50 | Autoimmune lymphoproliferative syndrome due to CTLA4 haploinsuffiency | UMLS:C4015214 |
| 51 | Autoimmune polyglandular syndrome type 1 | UMLS:C0085859 |
| 52 | Autosomal dominant hyper IgE syndrome | UMLS:C3489795 |
| 53 | Autosomal recessive candidiasis familial chronic mucocutaneous | UMLS:C3714992 |
| 54 | Autosomal recessive early-onset inflammatory bowel disease 28 | UMLS:C2751053 |
| 55 | Autosomal recessive early-onset inflammatory bowel disease 25 | UMLS:C2675508 |
| 56 | Barth syndrome | UMLS:C0574083 |
| 57 | Blau syndrome | UMLS:C1861303 |
| 58 | Bloom syndrome | UMLS:C0005859 |
| 59 | C1q deficiency | UMLS:C3150902 |
| 60 | Cartilage-hair hypoplasia | UMLS:C0220748 |
| 61 | CHARGE syndrome | UMLS:C0265354 |
| 62 | Chediak-Higashi syndrome | UMLS:C0007965 |
| 63 | Cherubism | UMLS:C0008029 |
| 64 | Combined immunodeficiency with skin granulomas | UMLS:C2673536 |
| 65 | Complement component 2 deficiency | UMLS:C3150275 |
| 66 | Complement component 8 deficiency type 1 | UMLS:C3151081 |
| 67 | Complement component 8 deficiency type 2 | UMLS:C3151080 |
| 68 | Cryoglobulinemic vasculitis | UMLS:C1852456 |
| 69 | Cyclic neutropenia | UMLS:C0221023 |
| 70 | Deficiency of interleukin-1 receptor antagonist | UMLS:C2748507 |
| 71 | Dendritic cell, monocyte, B lymphocyte, and natural killer lymphocyte deficiency | UMLS:C3280030 |
| 72 | Epidermodysplasia verruciformis | UMLS:C0014522 |
| 73 | Familial cold autoinflammatory syndrome | UMLS:C0343068 |
| 74 | Familial hemophagocytic lymphohistiocytosis | UMLS:C0272199 |
| 75 | Familial Mediterranean fever | UMLS:C0031069 |
| 76 | Felty’s syndrome | UMLS:C0015773 |
| 77 | Glycogen storage disease type 1B | UMLS:C0268146 |
| 78 | Griscelli syndrome type 2 | UMLS:C1868679 |
| 79 | Hepatic venoocclusive disease with immunodeficiency | UMLS:C1856128 |
| 80 | Hereditary folate malabsorption | UMLS:C0342705 |

| ID | Name | UMLS |
|---|---|---|
| 81 | Hermansky Pudlak syndrome 2 | UMLS:C1842362 |
| 82 | Hyper-IgD syndrome | UMLS:C0398691 |
| 83 | ICF syndrome | UMLS:C3279748 |
| 84 | IL12RB1 deficiency | UMLS:C4013949 |
| 85 | Immune dysfunction with T-cell inactivation due to calcium entry defect 1 | UMLS:C2748568 |
| 86 | Immunodeficiency with hyper IgM type 1 | UMLS:C0398689 |
| 87 | Immunodeficiency with hyper IgM type 2 | UMLS:C1720956 |
| 88 | Immunodeficiency with hyper IgM type 3 | UMLS:C1720957 |
| 89 | Immunodeficiency with hyper IgM type 4 | UMLS:C1842413 |
| 90 | Immunodeficiency with hyper IgM type 5 | UMLS:C1720958 |
| 91 | Immunodeficiency without anhidrotic ectodermal dysplasia | UMLS:C1845117 |
| 92 | Immune dysregulation, polyendocrinopathy and enteropathy X-linked | UMLS:C0342288 |
| 93 | Immunoglobulin A deficiency 2 | UMLS:C1836032 |
| 94 | Intestinal atresia multiple | UMLS:C0220744 |
| 95 | IRAK-4 deficiency | UMLS:C1843256 |
| 96 | Isolated growth hormone deficiency type 3 | UMLS:C0472813 |
| 97 | Leukocyte adhesion deficiency type 1 | UMLS:C0398738 |
| 98 | LRBA deficiency | UMLS:C3553512 |
| 99 | Majeed syndrome | UMLS:C1864997 |
| 100 | Melkersson-Rosenthal syndrome | UMLS:C0025235 |
| 101 | MHC class 1 deficiency | UMLS:C1858266 |
| 102 | MYD88 deficiency | UMLS:C2677092 |
| 103 | Netherton syndrome | UMLS:C0265962 |
| 104 | Neutrophil-specific granule deficiency | UMLS:C0398593 |
| 105 | Osteopetrosis autosomal recessive 7 | UMLS:C2676766 |
| 106 | Papillon Lefevre syndrome | UMLS:C0030360 |
| 107 | Activated PI3K-Delta Syndrome 2 | UMLS:C4014934 |
| 108 | Activated PI3K-Delta Syndrome | UMLS:C3714976 |
| 109 | Pearson syndrome | UMLS:C0342784 |
| 110 | PGM3-CDG | UMLS:C4014371 |
| 111 | Poikiloderma with neutropenia | UMLS:C1858723 |
| 112 | Pruritic urticarial papules plaques of pregnancy | UMLS:C0269680 |
| 113 | Purine nucleoside phosphorylase deficiency | UMLS:C0268125 |
| 114 | Pyogenic arthritis, pyoderma gangrenosum and acne | UMLS:C1858361 |
| 115 | Reticular dysgenesis | UMLS:C0272167 |
| 116 | Schimke immuno-osseous dysplasia | UMLS:C0877024 |
| 117 | Severe congenital neutropenia X-linked | UMLS:C1845987 |
| 118 | Short-limb skeletal dysplasia with severe combined immunodeficiency | UMLS:C1860168 |
| 119 | Shwachman-Diamond syndrome | UMLS:C0272170 |
| 120 | Spondyloenchondrodysplasia with immune dysregulation | UMLS:C1842763 |

| ID | Name | UMLS |
|---|---|---|
| 121 | T-cell immunodeficiency, congenital alopecia and nail dystrophy | UMLS:C1866426 |
| 122 | TARP syndrome | UMLS:C1839463 |
| 123 | Tumor necrosis factor receptor-associated periodic syndrome | UMLS:C1275126 |
| 124 | Vici syndrome | UMLS:C1855772 |
| 125 | WHIM syndrome | UMLS:C0472817 |
| 126 | Wiskott Aldrich syndrome | UMLS:C0043194 |
| 127 | Woods Black Norbury syndrome | UMLS:C1848144 |
| 128 | X-linked immunodeficiency with magnesium defect, Epstein-Barr virus infection and neoplasia | UMLS:C3275445 |
| 129 | X-linked lymphoproliferative syndrome | UMLS:C0549463 |
| 130 | ZAP-70 deficiency | UMLS:C2931299 |

#### Supplementary References

1. Himmelstein DS, et al. (2017) Systematic integration of biomedical knowledge prioritizes drugs for repurposing. *eLife* 6:e26726.
2. Drew K, et al. (2017) Integration of over 9,000 mass spectrometry experiments builds a global map of human protein complexes. *Mol Syst Biol* 13(6):932.
3. Giurgiu M, et al. (2019) Corum: the comprehensive resource of mammalian protein complexes–2019. *Nucleic Acids Res* 47(D1):D559–D563.
4. Türei D, et al. (2021) Integrated intra- and intercellular signaling knowledge for multicellular omics analysis. *Molecular Systems Biology* 17(3):e9923.
5. Pratt D, et al. (2015) Ndex, the network data exchange. *Cell Systems* 1(4):302–305.
6. Croft D, et al. (2014) The reactome pathway knowledgebase. *Nucleic Acids Res* 42(D1):D472–D477.
7. Köhler S, et al. (2021) The Human Phenotype Ontology in 2021. *Nucleic Acids Research* 49(D1):D1207–D1217.
8. Valdeolivas A, et al. (2018) Random walk with restart on multiplex and heterogeneous biological networks. *Bioinformatics* 35(3):497–505.
9. Wishart DS, et al. (2018) DrugBank 5.0: a major update to the DrugBank database for 2018. *Nucleic Acids Research* 46(D1):D1074–D1082.
10. Cheng F, Kovács IA, Barabási AL (2019) Network-based prediction of drug combinations. *Nature Communications* 10(1):1197.
11. Javierre BM, et al. (2016) Lineage-specific genome architecture links enhancers and non-coding disease variants to target gene promoters. *Cell* 167(5):1369–1384.e19.
12. Piñero J, et al. (2015) Disgenet: a discovery platform for the dynamical exploration of human diseases and their genes. *Database (Oxford)* 2015.
13. Piñero J, et al. (2020) The disgenet knowledge platform for disease genomics: 2019 update. *Nucleic Acids Res* 48(D1):D845–D855.
14. Brown AS, Patel CJ (2017) A standard database for drug repositioning. *Scientific Data* 4(1):170029.
15. Frankish A, et al. (2021) GENCODE 2021. *Nucleic Acids Research* 49(D1):D916–D923.
16. Cairns J, et al. (2016) Chicago: robust detection of dna looping interactions in capture hi-c data. *Genome Biology* 17(1):127.
17. Dixon JR, et al. (2012) Topological domains in mammalian genomes identified by analysis of chromatin interactions. *Nature* 485(7398):376–380.
18. Felix CA, et al. (1998) Association of CYP3A4 genotype with treatment-related leukemia. *Proceedings of the National Academy of Sciences* 95(22):13176–13181. Publisher: Proceedings of the National Academy of Sciences.
19. Kobayashi T, et al. (2010) Molecular and clinical analysis of RAF1 in Noonan syndrome and related disorders: dephosphorylation of serine 259 as the essential mechanism for mutant activation. *Human Mutation* 31(3):284–294.
20. Hasle H (2009) Malignant Diseases in Noonan Syndrome and Related Disorders. *Hormone Research* 72(Suppl. 2):8–14.
21. Oki T, et al. (2012) Aberrant expression of RasGRP1 cooperates with gain-of-function NOTCH1 mutations in T-cell leukemogenesis. *Leukemia* 26(5):1038–1045. Number: 5 Publisher: Nature Publishing Group.
22. Li H, et al. (2019) Knockdown of diacylglycerol kinase zeta (DGKZ) induces apoptosis and G2/M phase arrest in human acute myeloid leukemia HL-60 cells through MAPK/survivin/caspase pathway. *Die Pharmazie - An International Journal of Pharmaceutical Sciences* 74(7):418–422.
23. Trino S, et al. (2016) Inverse regulation of bridging integrator 1 and BCR-ABL1 in chronic myeloid leukemia. *Tumor Biology* 37(1):217–225.
24. Xu Y, Wertheim G, Morrissette JJD, Bagg A (2017) Braf kinase domain mutations in de novo acute myeloid leukemia with monocytic differentiation. *Leukemia & Lymphoma* 58(3):743–745. PMID: 27545333.
25. Zhang Y, et al. (2020) Role of RASA1 in cancer: A review and update (Review). *Oncology Reports* 44(6):2386–2396.
26. Lucena-Araujo AR, et al. (2011) High expression of AURKA and AURKB is associated with unfavorable cytogenetic abnormalities and high white blood cell count in patients with acute myeloid leukemia.

*Leukemia Research* 35(2):260–264.
27. Lee JW, et al. (2005) Mutational analysis of the ARAF gene in human cancers. *APMIS* 113(1):54–7. _eprint: https://onlinelibrary.wiley.com/doi/pdf/10.1111/j.1600-0463.2005.apm1130108.x.
28. Laverdière I, et al. (2018) Leukemic stem cell signatures identify novel therapeutics targeting acute myeloid leukemia. *Blood Cancer Journal* 8(6):1–16. Number: 6 Publisher: Nature Publishing Group.
29. Lethaby C, et al. (2007) Bisphosphonate Therapy for Reduced Bone Mineral Density During Treatment of Acute Lymphoblastic Leukemia in Childhood and Adolescence: A Report of Preliminary Experience. *Journal of Pediatric Hematology/Oncology* 29(9):613.
30. Auclair D, et al. (2007) Antitumor activity of sorafenib in FLT3-driven leukemic cells. *Leukemia* 21(3):439–445. Number: 3 Publisher: Nature Publishing Group.
31. Fan RF, et al. (2016) Zoledronic acid overcomes adriamycin resistance in acute myeloid leukemia cells by promoting apoptosis. *Molecular Medicine Reports* 14(6):5660–5666. Publisher: Spandidos Publications.
32. Fiedler W, et al. (2015) A phase I/II study of sunitinib and intensive chemotherapy in patients over 60 years of age with acute myeloid leukaemia and activating FLT3 mutations. *British Journal of Haematology* 169(5):694–700. _eprint: https://onlinelibrary.wiley.com/doi/pdf/10.1111/bjh.13353.
33. Kessler T, et al. (2019) Phase II clinical trial of pazopanib in patients with acute myeloid leukemia (AML), relapsed or refractory or at initial diagnosis without an intensive treatment option (PazoAML). *Annals of Hematology* 98(6):1393–1401.
34. Lainey E, et al. (2012) Erlotinib antagonizes ABC transporters in acute myeloid leukemia. *Cell Cycle* 11(21):4079–4092. Publisher: Taylor & Francis _eprint: https://doi.org/10.4161/cc.22382.
35. Martinez FO, Gordon S, Locati M, Mantovani A (2006) Transcriptional profiling of the human monocyte-to-macrophage differentiation and polarization: New molecules and patterns of gene expression. *The Journal of Immunology* 177(10):7303–7311.
